# Negative frequency-dependent selection and asymmetrical transformation stabilise multi-strain bacterial population structures

**DOI:** 10.1101/2020.06.05.118836

**Authors:** Gabrielle L. Harrow, John A. Lees, William P. Hanage, Marc Lipsitch, Jukka Corander, Caroline Colijn, Nicholas J. Croucher

## Abstract

*Streptococcus pneumoniae* can be split into multiple strains, each with a characteristic combination of core and accessory genome variation, able to co-circulate and compete within the same hosts. Previous analyses of epidemiological datasets suggested the short-term vaccine-associated dynamics of *S. pneumoniae* strains may be mediated through multi-locus negative frequency-dependent selection (NFDS), acting to maintain accessory loci at equilibrium frequencies. To test whether this model could explain how such multi-strain populations were generated, it was modified to incorporate recombination. The outputs of simulations featuring symmetrical recombination were compared with genomic data on locus frequencies and distributions between genotypes, pairwise genetic distances and tree shape. These demonstrated NFDS prevented the loss of variation through neutral drift, but generated unstructured populations of diverse isolates. Making recombination asymmetrical, favouring deletion of accessory loci over insertion, alongside multi-locus NFDS significantly improved the fit to genomic data. In a population at equilibrium, structuring into multiple strains was stable due to outbreeding depression, resulting from recombinants with reduced accessory genomes having lower fitness than their parental genotypes. As many bacteria inhibit the integration of insertions into their chromosomes, this combination of asymmetrical recombination and multi-locus NFDS may underlie the co-existence of strains within a single ecological niche.

## Introduction

One of the earliest applications of the term ‘strain’ to infectious disease referred to *“strains of lymph”*, pus containing cowpox or vaccinia viruses, that were propagated between individuals in smallpox vaccination programmes [1]. In the early 20^th^ century, strains of the bacterium *Streptococcus pneumoniae* (the pneumococcus) [2], an obligate commensal of the human nasopharynx, were differentiated by their ability to cause disease in a mouse model [3]. Similar variation in strains’ propensity to cause invasive diseases is observed in humans [4,5]. This underlies the effectiveness of the pneumococcal polysaccharide conjugate vaccines, which do not change nasopharyngeal carriage rates, but instead alter the bacterial population to reduce overall invasiveness [6]. Hence understanding pneumococcal strain dynamics is critical when designing interventions to reduce disease [5].

*S. pneumoniae*, like many other bacteria, was originally shown to cluster into genetically similar groups through electrophoretic analysis of polymorphic enzymes or restriction fragments [7–9]. Multi-locus sequence typing confirmed such clusters persisted despite ongoing exchange through recombination [10,11]. Population genomics has shown isolates within these clusters are similar in both core genome sequences and genome content [12,13]. The consistency of typing assignations by different methods suggests strains are meaningful biological entities [13,14]. Yet mirroring the broader problem of understanding bacterial speciation, co-existence of diverse genotypes in multi-strain populations (MSPs) has proved challenging to reproduce through evolutionary models [15–17].

Neutral models have sought to account for the persistence of MSPs in the absence of selected phenotypes. MSPs could be mixtures of ephemeral genotypes [18], with strains representing multiple sampling of local transmission chains, or microepidemics, within a diverse population [19]. This could account for the divergent strain compositions of cross-sectional samples from different countries [4,19,20]. Yet longitudinal sampling of individual populations [21,22] and phylodynamic reconstructions of strains’ evolutionary histories [4,23,24] concur that strains persist over decades. An alternative neutral explanation for high intraspecific diversity is allopatric diversification through “isolation by distance” [25,26]. However, this is unlikely to explain the diversity of *S. pneumoniae* strains, given their intercontinental distribution [4,23,24], and the similar frequencies of polymorphic loci in distant locations with different strain compositions [20]. This mixing is consistent with rapid migration between locations, which can facilitate population diversification if combined with repeated dissemination and local species-wide extinctions [27]. Yet *S. pneumoniae* is stably endemic worldwide [4,22].

Alternatively, transient separation of bacterial lineages may allow sufficient diversity to accumulate to inhibit homologous recombination, preventing subsequent convergence if they re-encounter one another [15,28]. However, neutral simulations predict even different streptococcal species would eventually converge through homologous recombination [15,29], as exemplified by previously-distinct *Campylobacter* species [30]. Correspondingly, interspecies recombinations between streptococci are readily detectable [31,32]. At the intraspecific level, there appear to be few mechanistic limitations to exchange of core loci between *S. pneumoniae* strains [33], with no evidence of increased frequencies of recombination between closely-related genotypes [34], and variable loci frequently diversifying through transformation [23,24,35].

Divergence may instead reflect bacterial ecology. In the ecotype model [36], two processes preserve strains’ distinctive genotypes. The first is “isolation by adaption” [37], with exchange between strains limited by their confinement to distinct [38], or even overlapping [39], niches. Yet with the possible exception of an atypical lineage [12,39], *S. pneumoniae* strains co-circulate in the same populations, while frequently directly competing with one another within individual hosts [40,41], implying their niches are insufficiently separate to prevent sequence exchange. The second process is selection against acquisition of locally-adaptive [42] or niche-specifying [43] loci. These are maladaptive in the recipient’s niche, but are individually insufficient to enable the recombinant to outcompete the donor genotype in its niche, assuming ecotypes’ adaptation to involve multiple loci [44,45]. This preserves MSPs by selecting against sequence exchange between different strains, a situation termed “outbreeding depression” [46], as recombinants’ fitness is typically reduced relative to the parental genotypes. However, *S. pneumoniae* strains have little private gene content [12,20,47], suggesting there are few stable niche-specifying loci. Such a pattern could instead indicate strains are less stably adapted to particular ecologies, with transiently-acquired mobile loci facilitating “recurrent invasion” [38] of different niches. However, the non-prophage component of *S. pneumoniae* accessory genomes is stable [12,13], with each possessing characteristic combinations of common loci [12], providing little scope for transient specialisation to “nano-niches” [43].

Strains may alternatively be adapted to different immunological niches, diversity in which results from variation in immune responses across a host population [48–50]. Variable antigens are likely to be under negative frequency-dependent selection (NFDS), which results from a phenotype conferring its greatest benefit to an individual when it is rare in the population [51], as epitopes are more frequently recognised by adaptive immunity when they are common [52,53]. If multiple strongly immunogenic antigens are simultaneously under NFDS, then strains are predicted to emerge from a freely-recombining population through selection of discordant antigen combinations that minimise cross-strain immunity [54]. MSPs are thereby maintained by outbreeding depression resulting from recombinants being recognised by cross-immunity induced by either parental strain [55]. However, the distribution of antigen variants is not strongly discordant across *S. pneumoniae* populations [56], which may reflect immunity induced by *S. pneumoniae* colonisation only weakly protecting against reinfection by the same antigen profile [57,58], or the immunity induced by exposure to one strain recognising much of the species-wide diversity of even variable antigens [59].

This model also predicts the structuring of MSPs by immunity should change with hosts’ contact network [60], which is not apparent from comparisons of Western *S. pneumoniae* populations with those from a more isolated refugee camp [20]. Strong immune selection is also predicted to drive similar levels of diversity at each antigenic locus [61], whereas *S. pneumoniae* antigens exhibit very different diversity levels within- and between-strains [22–24,56].

Discordant antigen models have been extended to incorporate virulence [21] and ecotype-style metabolic adaptation to niches through particular combinations of core gene alleles [62]. More generally, strains may emerge as particular combinations of co-evolving loci [46]. The disruption of advantageous epistatic interactions contributing to each strain’s fitness can cause sufficient outbreeding depression to maintain MSPs [46]. However, the polymorphic loci that define strains are individually found in many combinations across the species [12,47], and genome-wide epistasis analyses suggest co-evolutionary associations are focussed on a few key loci [63,64]. Hence none of the models of MSPs conceived prior to routine population genomics studies can be easily reconciled with such datasets, and there is an opportunity to use this information to improve our understanding of bacterial evolution [27,65].

One model developed using multiple collections of genomic data assumed all intermediate-frequency accessory loci (*i.e*., those present at between 5% and 95% prevalence in the population) were maintained at “equilibrium frequencies” by NFDS acting on multiple phenotypes (the multi-locus NFDS model [20]). This framework was able to replicate the short-term post-vaccine dynamics of *S. pneumoniae* MSPs [20,66]. The multi-locus NFDS model differs from its antecedents in that genotypes compete within an homogeneous niche across multiple loci, each of which independently contributes to a genotype’s overall fitness. Yet the model did not feature recombination, and therefore whether it was consistent with the emergence and persistence of MSPs was unclear.

The majority of pneumococcal intermediate-frequency accessory loci are not autonomously mobile [20], and therefore their transfer primarily occurs through homologous recombination, which is frequent in the naturally transformable pneumococcus [67]. While core loci are typically modelled as being symmetrically exchanged (*i.e*., all alleles transfer at equal rates per donor), transformation with accessory loci is asymmetric. This is because any multi-gene accessory locus can be deleted, through recombination between a recipient encoding the locus and a donor lacking it, substantially more efficiently than it can be inserted through the reciprocal recombination [68]. Hence the multi-locus NFDS model was modified to incorporate different modes of recombination affecting the core and accessory genomes, and simulations were run to test whether MSPs could be maintained over longer timescales.

## Materials and Methods

### Structure of the model

The previously described multi-locus NFDS model [20] was constructed using a Wright-Fisher framework. At time *t*, each isolate *i* contributed a Poisson-distributed number of progeny, *X_i,t_*, to the next generation. The total number of *i* in the population at *t*, *N_t_*, was constrained by density-dependent selection, parameterised by the carrying capacity, *κ*. Each individual *i* in the model encoded *L* biallelic accessory loci, each denoted *l*, as encoded by the binary matrix *g_i,l_*. If *g_i,l_* = 1, *l* was present in *i*; if *g_i,l_* = 0, it was absent. NFDS was incorporated through comparing the instantaneous frequency of *l* at time *t, f_l,t_*, with its equilibrium frequency, *e_l_*. Therefore, the fitness of *i* was dependent on π_*i,t*_, quantifying the extent to which a genotype encoded accessory loci at instantaneous frequencies that were above, or below, their equilbrium frequencies:

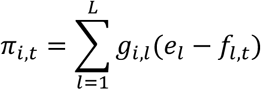

The effect on *X_i,t_* was mediated by the strength of NFDS, as parameterised by *σ_f_*:

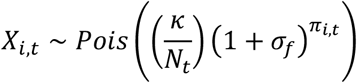

The model was modified in three ways to enable evolutionary simulations to be carried out. The first was to enable exchange of *l* between *i* through transformation. This was parameterized by three variables:

- Transformation rate, τ: the per-timestep probability of *i* being a recipient, *r*, in an exchange of DNA through transformation
- Proportion of genotype affected by transformation, ρ: the per-transformation probability of each locus *l* being exchanged by recombination
- Transformation asymmetry, ϕ: the per-recombination probability of *l* being acquired if it was present in a donor, *d*, but not *r*

Each *i* in the extant population underwent transformation with a probability τ in each generation. For each transformation, a single *d* was selected from the extant population at random. Each *l* in *g_r,l_* underwent recombination with probability ρ. If *g_r,l_* = *g_d,l_, g_r,l_* was unmodified. If *g_r,l_* = 1 and *g_d,l_* = 0, then *l* was deleted (*g_r,l_* set to 0). If *g_r,l_* = 0 and *g_d,l_* = 1, *l* was inserted with probability ϕ, else *g_r,l_* remained unmodified. The *f_l,t_* values were recalculated following transformation, prior to the calculation of *X_i,t_*, to enable NFDS to reflect the altered gene frequencies.

The second modification was the incorporation of *S* neutral loci, corresponding to core genome single nucleotide polymorphisms (SNPs; each denoted *s*), into the genotype of *i*. These were also biallelic, and encoded in a matrix *c_i,s_*. These were not under NFDS, and for each pair of *d* and *r*, each *s* underwent recombination with the same probability, ρ. However, transformation was always symmetrical (ϕ = 1) for *s*.

The third modification was the incorporation of a time-varying pool of migrants to enable the effects of allopatric diversification to be tested, without interfering with the temporal dynamics of the evolutionary simulations. At time *t*, *N_m,t_* isolates were imported into the simulated population at a rate determined by the parameter *m*:

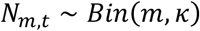

Hence the reproduction function was changed to:

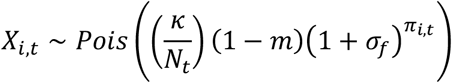

Each migrant pool was extracted from simulations run for discrete intervals of *j*_1_, *j*_2_… *j_k_* generations. While *j_k_* ≤ *t* < *j*_*k*+1_, the migrant pool consisted of *i* from simulations run for *j_k_* generations. Each *i* in the pool was equally likely of being randomly drawn to contribute to the *N_m,t_* imported isolates.

### Input data for simulations

The input data were the 616 genomes from the Massachusetts *S. pneumoniae* collection [22], processed as described previously [20]. Only the 1,090 *L* present at intermediate frequencies (*i.e*. between 5% and 95%) in the peri-vaccination population were included as being under NFDS in the simulations. For simulating the evolution of core genome variation, 1,090 biallelic core genome single nucleotide polymorphism (SNPs), in which the minor allele frequency was above 5%, were randomly selected from the genomic data to be included in the simulations. Hence *L* and *S* were equal in magnitude, to simplify the comparison of pairwise distances calculated from *g_i,l_* and *c_i,s_*. At each *s*, the allele denoted as ‘0’ matched that within the sequence of *S. pneumoniae* ATCC 700669 [69], such that the SNP frequency at *t*(*f_s,t_*) was that of the alternative allele. This ensured *f_l,t_* and *f_s,t_* were distributed over similar ranges.

### Calculation of parameter values

Each generation corresponded to a month, and κ was 10^5^, to enable parameters estimated from previous model fitting to the Massachusetts *S. pneumoniae* population to be used with these simulations [20]. The *e_l_* were also calculated from the peri-vaccination sample from these data [20,22].

NFDS acted homogeneously on each accessory locus. The value *σ_f_* used in multi-locus NFDS simulations (0.0356 month^-1^) was calculated from the fitted model parameters as a weighted mean of the strong (*σ_f_* = 0.1363 month^-1^) and weak (*σ_w_* = 0.0023 month^-1^) NFDS strengths, according to the fraction (*p_f_* = 0.2483) subject to *σ_f_* [20]. Simulations were also run with ‘weak NFDS’, using only the latter value (*σ_w_* = 0.0023 month^-1^).

The rate of inter-strain transformation was based on the best-fitting homogeneous rate recombination model (model 2: null model with over-dispersion) from a modelling study of divergence through recombination in *S. pneumoniae* strains [70]. Transformations detectable through exchanging sequence variation between strains were estimated to occur at a mean rate, τ, of 0.21 y^-1^ (τ = 0.0175 month^-1^) and span 6.4 kb of the genome. This rate does not include within-strain transformations, which are unlikely to be detectable through sequence divergence. Additionally, as these values are inferred from isolate collections, they correspond to the post-selection rate, and therefore will underestimate the actual transformation rate.

Across the 616 genomes in the Massachusetts dataset, isolates encoded a mean of 309 intermediate-frequency loci. As an *S. pneumoniae* genome contains ~2,000 genes [22], this implies intermediate-frequency loci comprise ~15% of the genome. Therefore the monthly whole genome transformation rate (τ = 0.0175 month^-1^) was scaled to represent the rate with which transformation would affect only the intermediate-frequency loci, assuming an homogeneous distribution of transformation events (τ = 0.002625 month^-1^).

The typical transformation event size (6.4 kb [70]) would be consistent with the acquisition or deletion of a 5 kb accessory locus, given that transformation events affecting accessory loci typically include two flanking homologous arms of 0.75-1 kb [68]. Such an homologous recombination would correspond to approximately five intermediate-frequency genes, given each *S. pneumoniae* gene has a length of ~1 kb [69]. As each isolate encoded approximately 300 accessory loci, the proportion of loci affected during an exchange through transformation (ρ) was set to 0.0167, such that the expected number of loci present in the donor and recipient potentially affected by transformation was approximately five. Given this size of accessory locus and the experimentally-determined relationship between the efficacy of insertion relative to SNP transfer [68], the magnitude of transformational asymmetry (ϕ) was estimated to be 0.05.

The product τρ, representing the mean rate at which transformation affects each *S. pneumoniae* accessory locus, dictates the timescale over which the effects of recombination are detectable. The calculated value of 4.38×10^-5^ month^-1^ indicated each locus would only be expected to be affected by transformation in a given isolate once every ~2,000 years. Therefore simulations were run for 60,000 generations (equivalent to ~5,000 years).

Transformation appears to be a saltational process, with substantial inter-strain exchanges occurring infrequently [70]. These rare, but extensive, recombination events may play an important role in the emergence of strains [71]. To model this, simulations were run in which τ was reduced five-fold (τ = 0.000525 month^-1^), and ρ increased five-fold (0.0835), such that the mean transformation rate per locus (τρ) remained constant.

To include the effects of a geographically-structured population, simulations were also run with inward migration from ten external populations. High rates of migration would be expected to cause all communities within a metapopulation to homogenise [72]. Hence *m* was set at 10^-5^, such that under neutral evolution it would be expected that the majority of isolates in a population at the end of the simulations would not have been imported from other sources (i.e., (1-*m*)^60000^ > 0.5).

### Details of simulations

The *S. pneumoniae* population was initialised at size κ, as a random mixture of the 616 genotypes sequenced in a genomic study in Massachusetts [22]. For each combination of selection (neutral; multi-locus NFDS; weak multi-locus NFDS) and transformation (none; symmetrical; asymmetrical; saltational symmetrical, and saltational asymmetrical), 100 independent stochastic simulations were run for 60,000 timesteps (corresponding to 5,000 years; parameter combinations are summarised in Table 1).

**Table 1:**
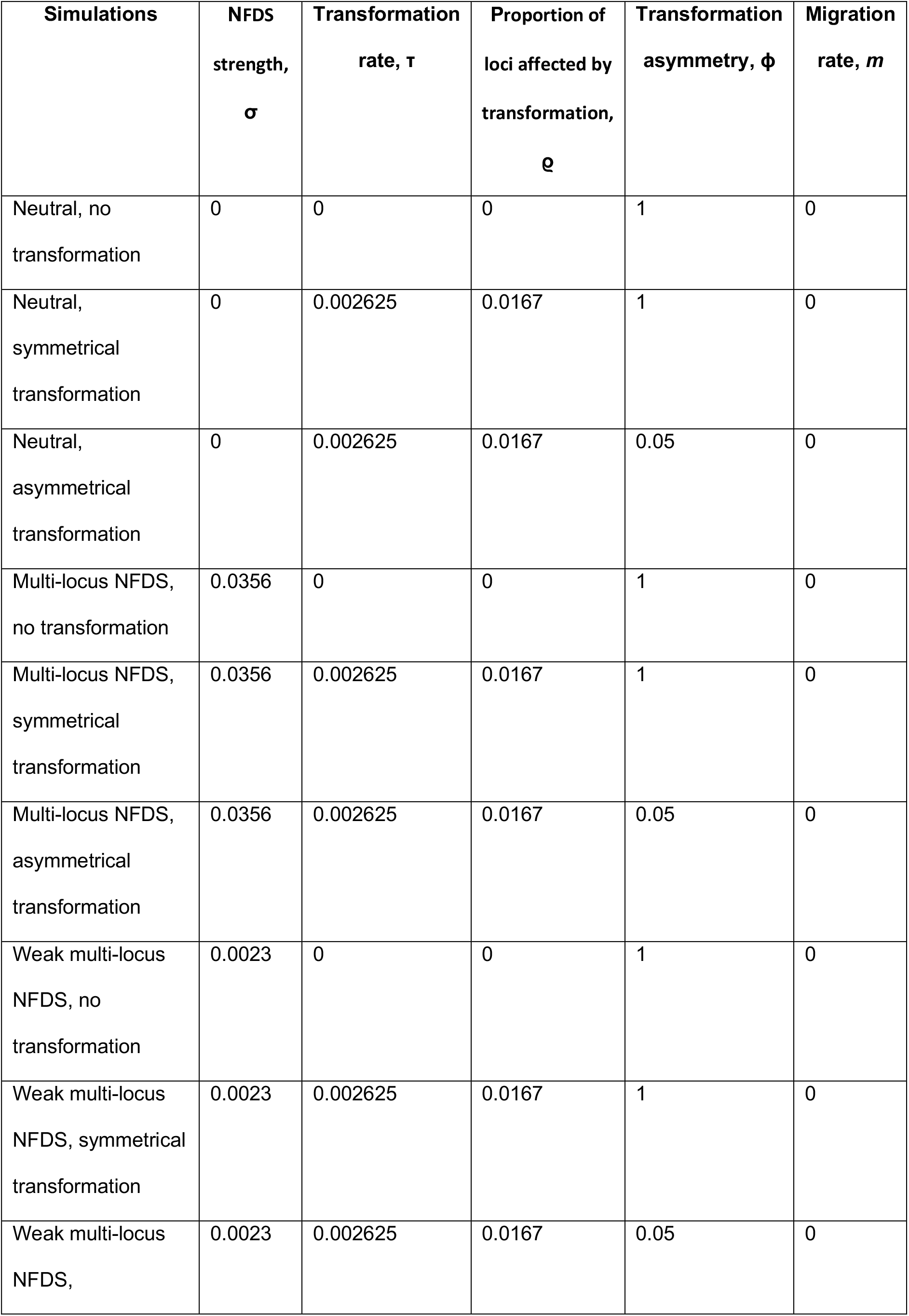
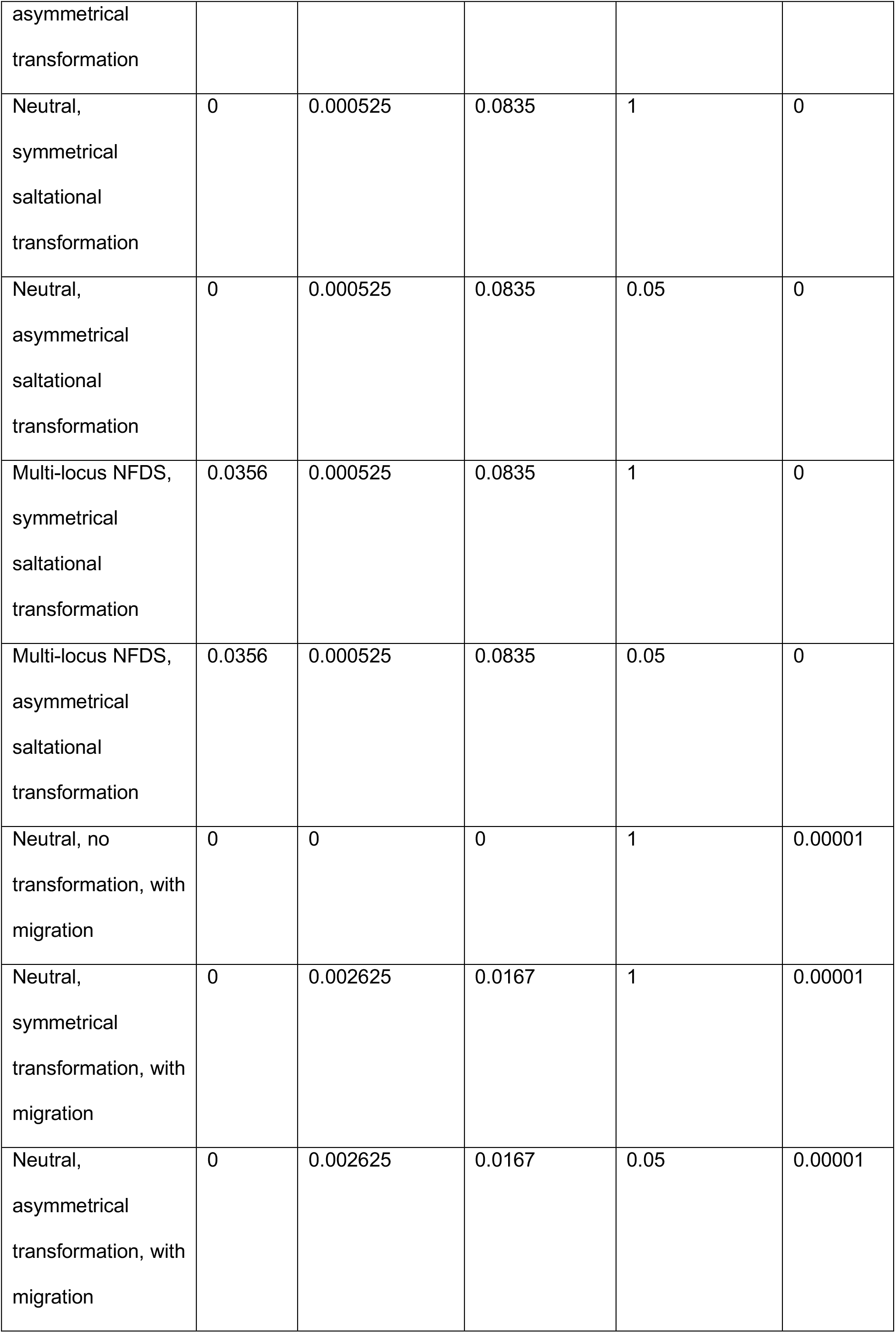
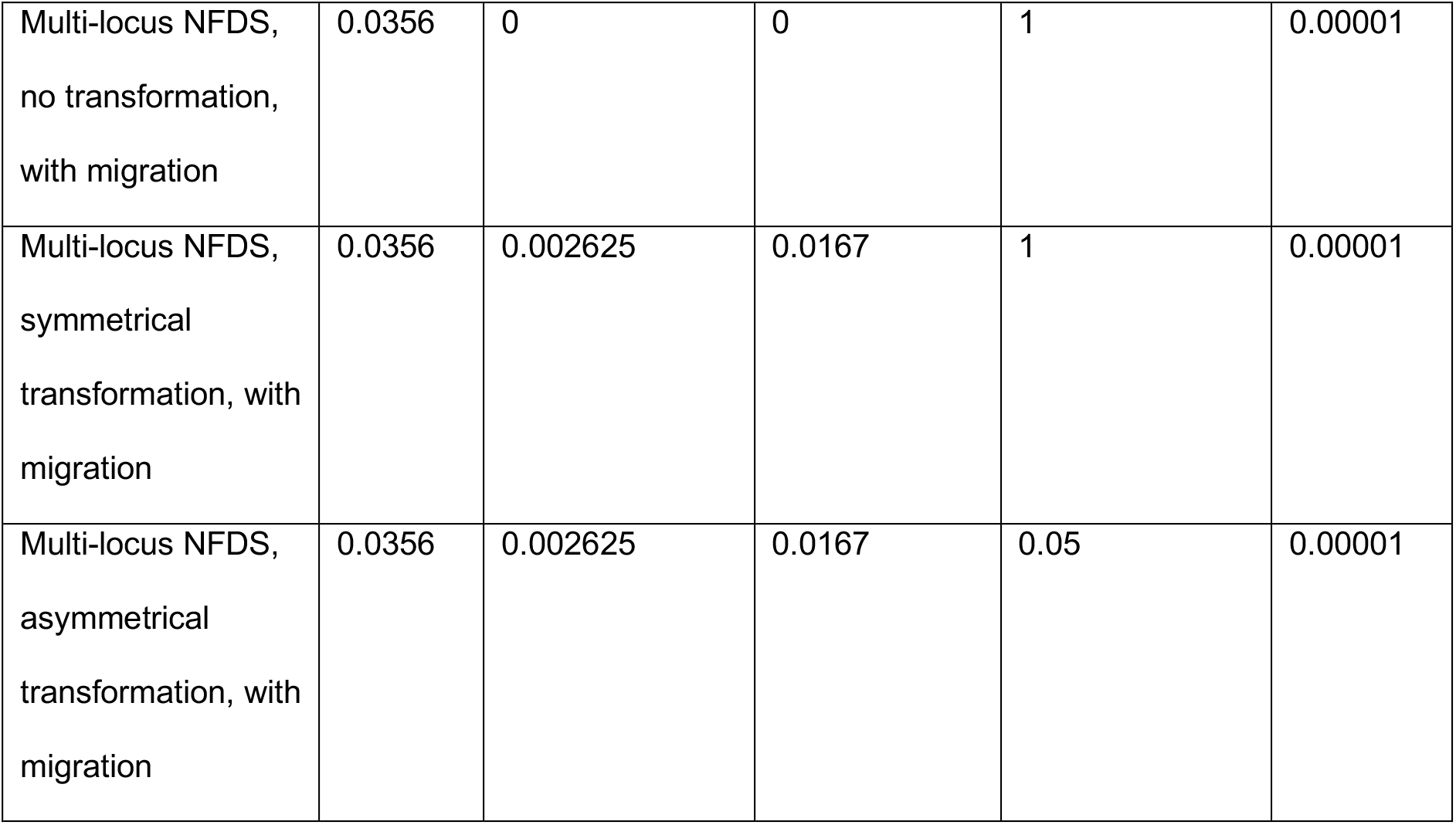
Parameter values for each set of simulations. In each simulation, κ = 10^5^.

To ensure the evolutionary dynamics of incoming migrant genotypes were synchronised with the simulated population, ten independent replicate source populations were generated prior to the reported simulations. Each of these was run as a series of 600 timestep simulations, up to a maximum of 60,000 timesteps, without migration. At the end of each 600 timestep interval, 5,000 genotypes were randomly sampled from the final generation (without consideration of the population structure), and used to generate a population of size κ to initiate the next phase. Once these serial simulations were complete, ten isolates were randomly drawn from each sampling timepoint in each of the ten replicates. These were combined to generate a pool of 100 migrants for each of the *j*_1_, *j*_2_… *j_k_* intervals for each set of equivalently-parameterised simulations featuring migration (see above).

### Statistical analyses

At the end of each simulation, 616 isolates were randomly sampled from the final timestep, to match the input data. Analyses and visualisations used R [73] with Tidyverse [74] and ggpubr [75] packages. Distance calculations were performed with rdist [76] and distributions analysed with propagate [77]. Neighbour-joining trees were constructed using ape [78] and visualised with ggtree [79]. Pybus and Harvey’s γ [80] was calculated using ape after trees were converted to be ultrametric using phytools [81]. Strains were defined using the genetic distance threshold indicated in figure 4 [13] by constructing networks using igraph [82]. Permutations of gene presence and absence were conducted with vegan [83].

The code used for simulations, and analysis of outputs, is available from https://github.com/nickjcroucher/multilocusNFDS/.

## Results

### Multi-locus NFDS prevents transformation-mediated decay of the accessory genome

One hundred simulations were run for each of six parameter combinations, corresponding to the absence of transformation, symmetrical transformation, or asymmetrical transformation, either under neutrality or multi-locus NFDS (see Methods). The frequencies of accessory loci at simulation endpoints (*f*_*l*,60000_) were compared to the starting frequencies (*f*_*l*,0_; Fig. 1a). In neutral simulations in which transformation was absent or symmetrical, *f*_*l*,0_ and mean *f*_*l*,60000_ correlated significantly. Yet *f*_*l*,60000_ varied extensively across replicate simulations through drift, with diversity often lost at loci with extreme *f*_*l*,0_ values, consistent with initial allele frequency determining probability of fixation [84]. However, asymmetrical transformation reduced mean *f*_*l*,60000_ and increased the probability of the null, or empty, allele fixing in the population [68]. In such simulations, there was a disproportionate loss of rare accessory loci, resulting from the high prevalence of recombination donors lacking the locus able to cause deletion in recipients originally possessing the locus. This reduced the overall number of accessory loci per genome (Fig. 2). Hence, under neutrality, asymmetric transformation accelerates the loss of accessory loci through drift.

**Figure 1:**
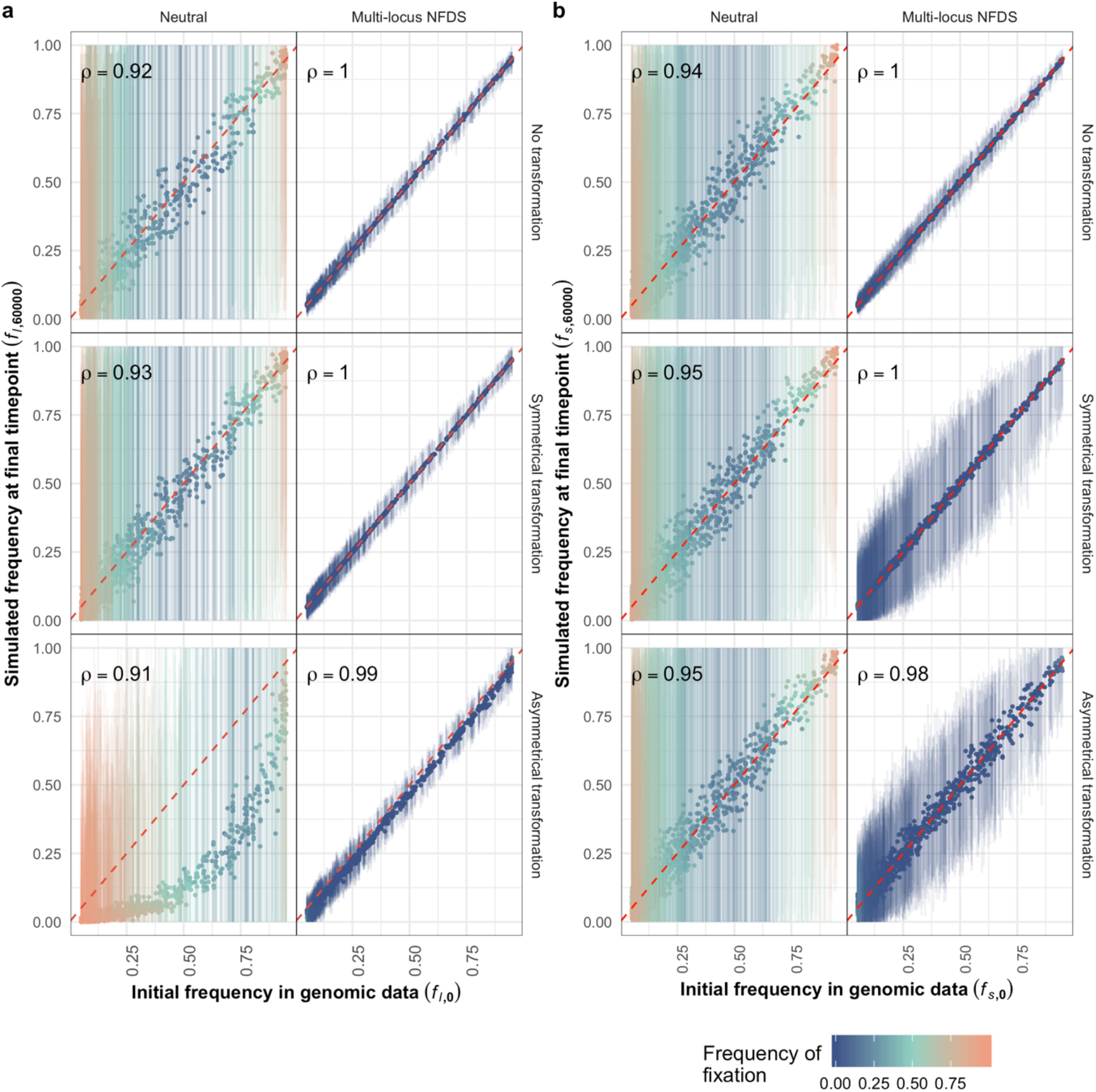
Scatterplots comparing the frequency of alleles at the initial timepoint in the genomic data to their frequency in the final simulation timepoint (*N* = 616 isolates sampled from each simulation). Each point (*L* or *S* = 1,090 in each panel) represents the mean, and the vertical lines show the range, from 100 replicate simulations. These are coloured by the frequency with which an allele fixed at the corresponding polymorphic locus (*i.e*. the displayed allele’s final frequency was zero or one in an individual simulation). Spearman’s correlation statistic (**ρ**) is shown in each panel; all *p* values were <10^-10^. **a** Frequency of accessory loci, in which the alleles correspond to the presence of intermediate-frequency genes. The initial frequencies correspond to the equilibrium frequencies in the model. **b** Allele frequencies at single nucleotide polymorphism sites.

The loss of variation through both drift and asymmetrical transformation was prevented by multi-locus NFDS, which strengthened the correlation between *f*_*l*,0_ and mean *f*_*l*,60000_, and reduced the variation in *f*_*l*, 60000_ across replicates (Fig. 1a). Correspondingly, the mean number of accessory loci per isolate was maintained close to that observed in the genomic data (Fig. 2). These simulations also replicated the positive skew of the sequenced genomes’ accessory locus content, unless transformation was symmetrical. Symmetrical exchange instead homogenised genome characteristics, under neutrality and multi-locus NFDS, reducing both the variance and skew of the distribution of accessory loci per isolate. Excepting those simulations in which isolates’ accessory locus content was preserved by a lack of transformation, the overall distribution of locus content was best reproduced by the combination of multi-locus NFDS and asymmetrical transformation (Fig. S1).

**Figure 2:**
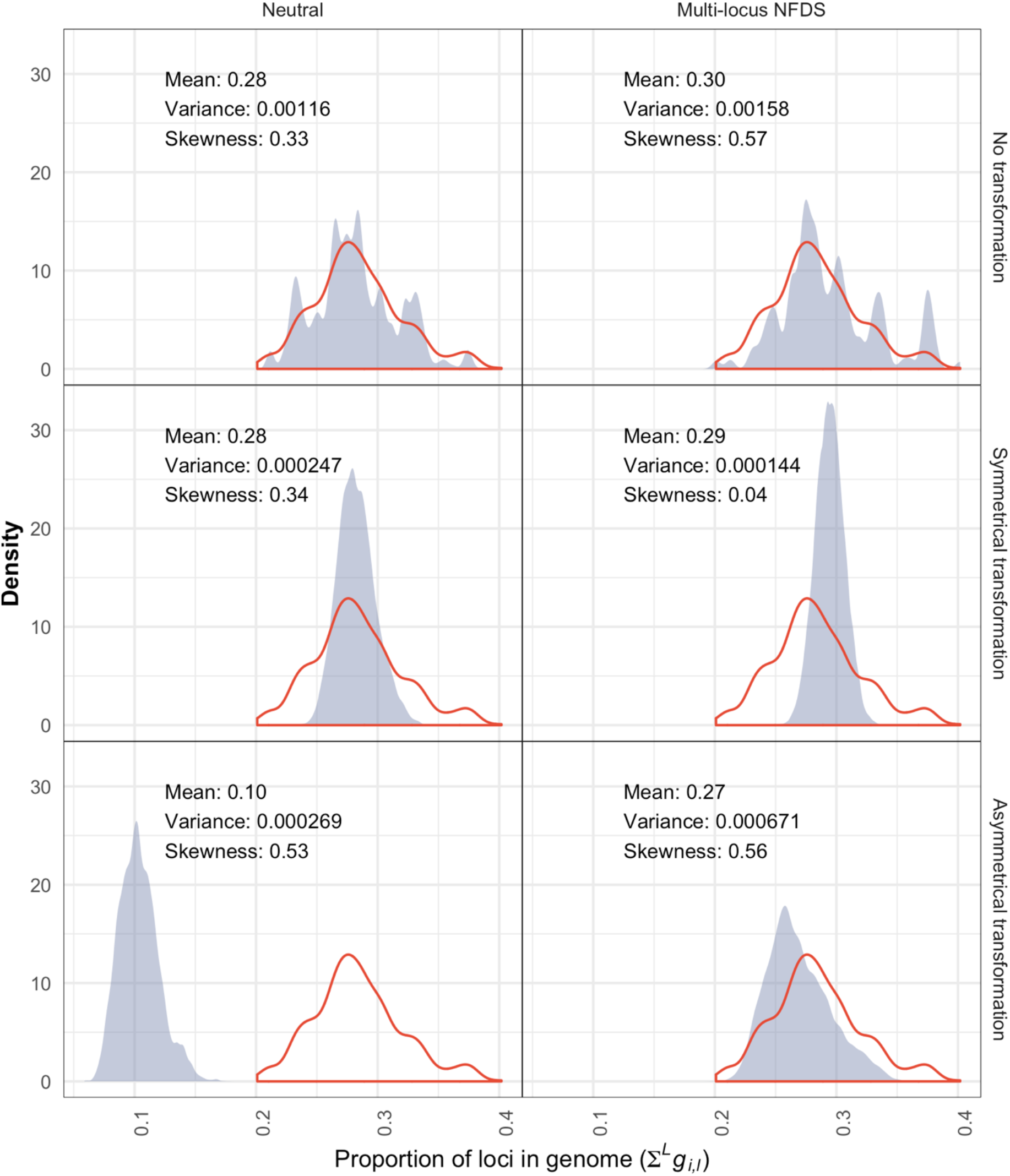
Density plots comparing the variation in gene content in the genome data and at the final timepoint of simulations. The horizontal axis represents to the proportion of intermediate-frequency loci (*L* = 1,090) present in isolates (calculated as *Σ^L^g_i,l_* for each *i*). The red outline shows the distribution from the 616 genomes in the original dataset (mean: 0.28, variance: 0.00132, skewness: 0.45). In each panel, the blue shading and displayed statistics describe the overall distribution (*N* = 61,600) from 616 isolates sampled from the final timepoint of 100 replicate simulations.

Further simulations tested the robustness of these conclusions to plausible alterations to parameters (Table 1). When transformation was saltational, such that exchanges were more extensive but less frequent, few differences were observed (Fig. S2). Neutral simulations featuring inward migration from external populations (see Methods) exhibited less variation in *f*_*l*,60000_ across replicates, as importation of genotypes mitigated some loss of diversity through drift (Fig. S3). However, reducing the strength of multi-locus NFDS increased the variation in *f*_*l*,60000_ (Fig. S4), and failed to prevent asymmetrical transformation reducing genome content (Fig. S5). Hence there is a threshold NFDS strength required to prevent loci being lost through asymmetrical transformation.

### Multi-locus NFDS stabilises frequencies of unselected polymorphisms

SNPs were also included in the simulations (see Methods). These always evolved neutrally, and, if transformation occurred, were exchanged symmetrically. In neutral simulations, the mean final SNP frequencies (*f_s,60000_*), and probabilities of fixation, were similar to those of the accessory loci when transformation was absent or symmetrical (Fig. 1b). Multi-locus NFDS indirectly stabilised the SNP frequencies, despite only acting directly on the accessory loci. This effect was greatest in the absence of transformation, such that all accessory and SNP loci maintained the linkage embedded in the original population. Although mean *f*_*s*,60000_ was similarly strongly correlated with *f*_*s*,0_ when transformation was symmetrical or asymmetrical, there was greater variation between replicate simulations, likely reflecting the weakened linkage with the selected accessory loci.

This correlation strengthened if asymmetrical transformation were also saltational (Fig. S2), but inward migration had little effect (Fig. S3). Weaker NFDS increased the variation in *f*_*s*,60000_ between replicates across all non-neutral simulations (Fig. S4). Therefore, the correlation between *f*_*s*,0_ and *f*_*s*,60000_ was higher when multi-locus NFDS was stronger.

### Pairwise distance distributions are shaped by selection and recombination

The variation in gene content across the population was described by calculating the pairwise binary Jaccard distances between all genotypes sampled at the simulation endpoints, based on the intermediate-frequency accessory loci they encoded. In the genomic data, there is a single mode of between-strain distances, and a tail of shorter within-strain distances that make the distribution negatively skewed (Fig. 3a). In the neutral simulations, the mean pairwise distance, and negative skew, are reduced through two mechanisms. Firstly, the loss of genotypes through drift means the within-strain peak increases in prominence relative to the between-strain peak, as the population simplifies. Secondly, transformation drives convergence between genotypes, such that the range of pairwise distances is decreased [13,85,86]. The effect is least pronounced for asymmetrical transformation, as the number of accessory loci per genome (to which the Jaccard distance is inversely proportional) is decreased across the population (Fig. 2).

**Figure 3:**
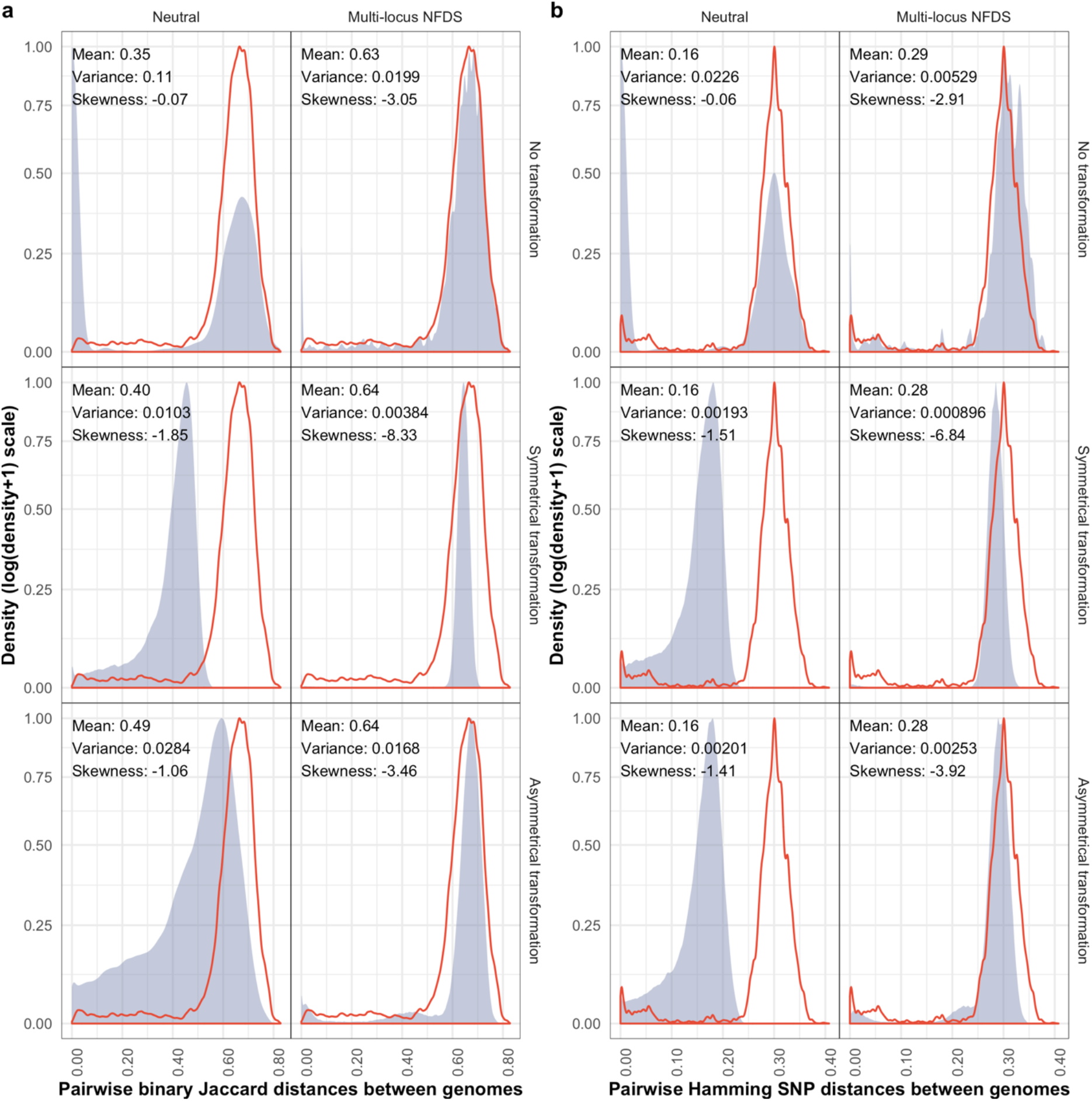
Density plots comparing the distribution of pairwise genetic distances between isolates in the genome data and at the final timepoint of simulations. The red outline shows the distribution from the genome data (*N* = 189,420). The blue shading and displayed statistics in each panel describe the overall distribution of a random 2% sample of the equivalent distances calculated from the final timepoint of 100 replicate simulations (overall *N* ≅ 378,840). **a** Pairwise binary Jaccard distances calculated from isolates’ accessory loci compared with genomic data (mean: 0.63, variance: 0.015, skewness: −2.91). **b** Pairwise Hamming distances calculated from single nucleotide polymorphisms compared with genomic data (mean: 0.29, variance: 0.0043, skewness: −3.06).

Multi-locus NFDS maintains the position of the between-strain mode, regardless of transformation. However, the tail of within-strain distances was only maintained when transformation was absent or asymmetrical. Symmetrical transformation instead homogenised the distance distribution, resulting in a unimodal distribution with low variance. Consequently, multi-locus NFDS simulations consistently reproduced the observed pairwise distance distribution more accurately than neutral equivalents, and simulations featuring symmetrical transformation were the least accurate (Fig. S6).

The pairwise Hamming distances were also calculated from the SNP alleles encoded by the simulated genotypes (Fig. 3b). In the genomic data, these again have a between-strain peak, and a tail of shorter within-strain distances. Each of these simulation outputs was similar to the corresponding set of accessory genome distances (Fig. S7), with the exception of neutral simulations featuring asymmetrical transformation, in which SNPs were symmetrically exchanged. Hence all the neutral simulations had a mean pairwise Hamming distance approximately half that calculated from the genomic data. In all multi-locus NFDS simulations, the mean divergence was maintained near its observed value. However, symmetrical transformation again homogenised the pairwise distance distribution, and hence the tail of within-strain small SNP distances was only evident if transformation of accessory loci were absent or asymmetric.

Saltational transformation had little effect on the pairwise distances (Fig. S8). Inward migration altered the output from neutral simulations, increasing the mean pairwise distances between genomes through the import of strains, which mitigated the loss of diversity through drift (Fig. S9). Weaker multi-locus NFDS was less effective at preventing transformation driving convergence in the core genome, but still maintained the observed accessory genome distances (Fig. S10). Hence multi-locus NFDS preserved between-strain variation, and if transformation were absent or asymmetrical, within-strain distances were also replicated.

### Multi-locus NFDS and asymmetrical transformation stabilise MSPs

MSPs can be characterised by a positively correlated, discontinuous distribution of accessory and core genome pairwise distances [12,13]. All parameter sets generated positive correlations in pairwise distances, but the discontinuity separating within- and between strain distances was less consistently preserved (Fig. 4). This is a consequence of reduced diversity in neutral simulations. Without transformation, the plot became sparse, with most comparisons between identical genotypes at the origin of the graph. With transformation, the convergence between genotypes resulted in a continuous distribution of points, which no longer spanned the observed range of genetic distances. By contrast, simulations with multi-locus NFDS maintained the between-strain distances near their observed position. However, when transformation was symmetrical, there was an absence of within-strain points. Therefore, multi-locus NFDS only generated MSPs when transformation was asymmetric or absent.

**Figure 4:**
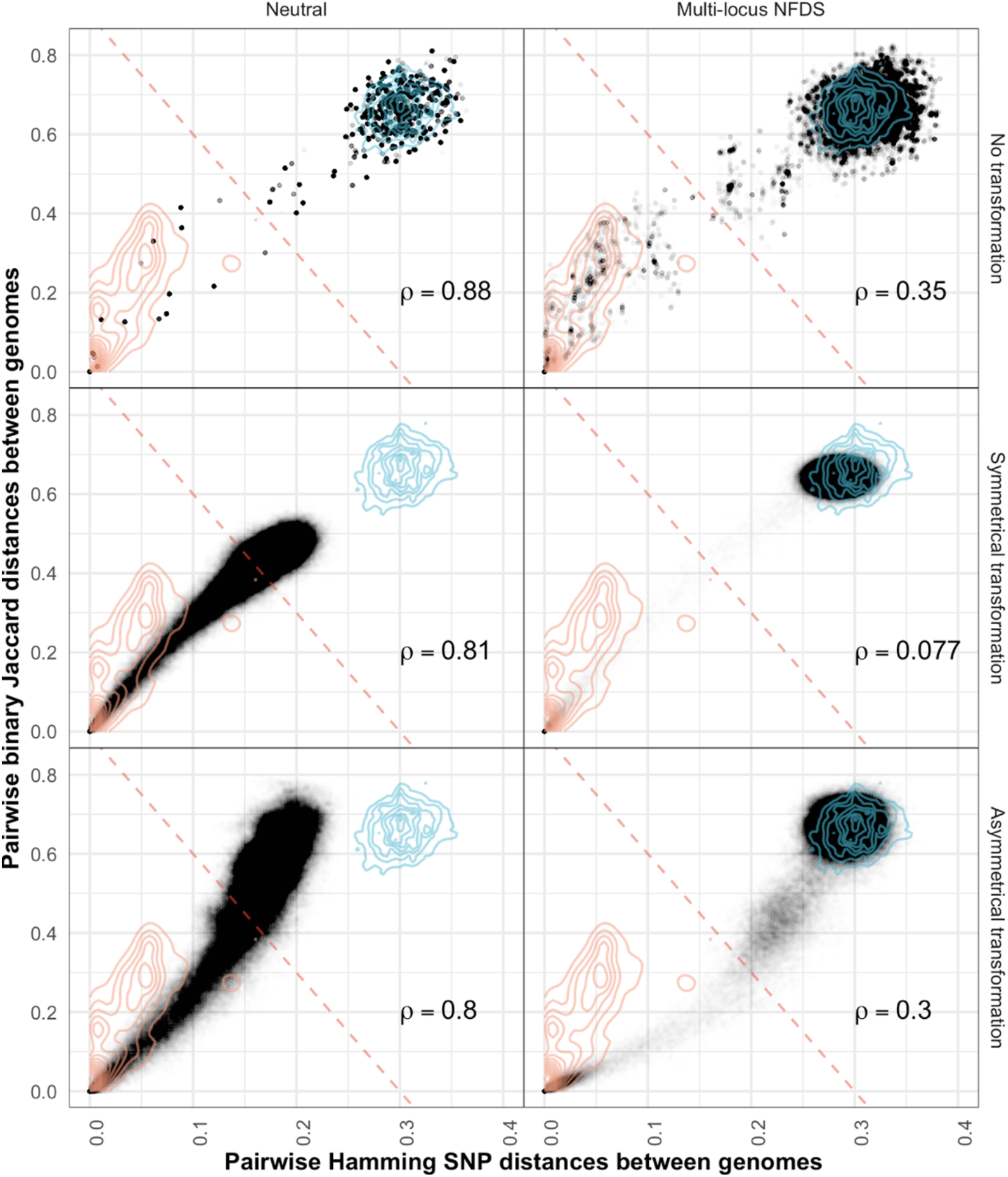
Scatterplots comparing the distributions of pairwise genetic distances between isolates at the final timepoint of simulations. Each panel combines the data from the equivalent panels in Fig. 3 (*N* ≅ 378,840). Each point is a single comparison between isolates, with the horizontal axis representing divergence in core genome single nucleotide polymorphisms, and the vertical axis representing divergence in accessory loci. The red diagonal is a threshold distinguishing within- and between-strain distances (Fig. S11). The contours describe the distribution of within- and between-strain pairwise distances in the genomic data (orange and blue lines, respectively; *N* = 189,420). Spearman’s correlation statistic (**ρ**) is shown in each panel; all *p* values were <10^-10^.

A diagonal boundary can be used to separate within- and between-strain distances in the genomic (Fig. S11) and simulated (Fig. 4) data, and thereby define strains using the approach implemented by PopPUNK [13]. A strain rank-frequency plot demonstrates *S. pneumoniae* populations typically consist of a few common strains, and a tail of many rare genotypes [4] (Fig. 5). Neutral simulations consistently generated populations containing few dominant strains, as rare genotypes were lost through drift, and transformation drove the merging of previously-distinct strains [28,87]. By contrast, simulations combining multi-locus NFDS with symmetrical transformation output populations in which all strains were rare, consistent with the high genetic diversity of the population. A mixture of common and rare strains was only observed when transformation was asymmetric or absent.

**Figure 5:**
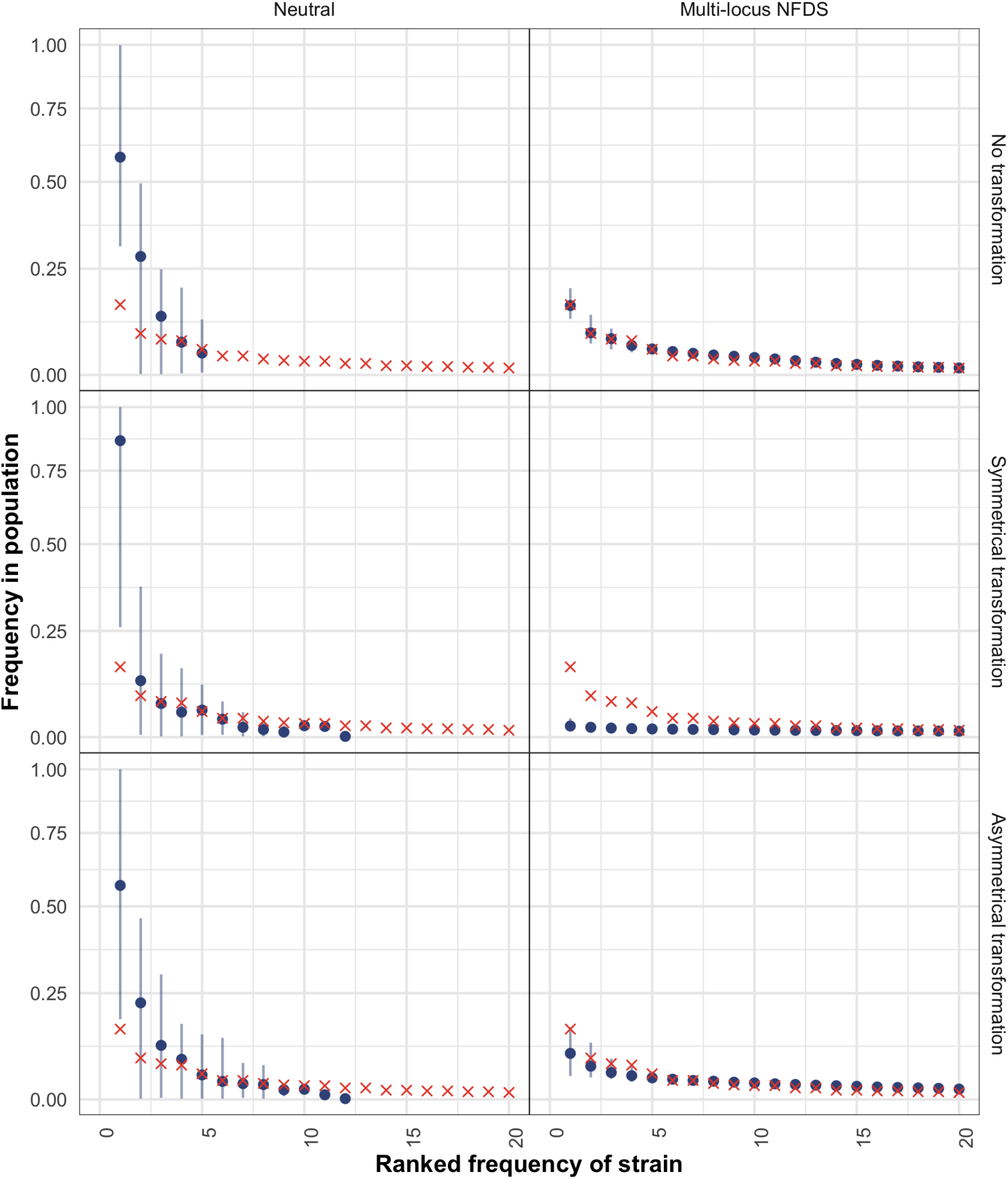
Scatterplots comparing the rank-frequency distributions of strains in the overall set of 616 genomes (red crosses), and those from samples of 616 isolates from the final timepoint of 100 replicate simulations. The blue points show the mean, and the vertical lines show the range, of strain frequencies for each observed rank across replicate simulations.

Saltational recombination had little effect on the MSPs, except that the output of simulations featuring multi-locus NFDS and asymmetric transformation more closely mirrored the observed distribution (Fig. S12, S13). Migration instead improved the fit of neutral simulations, through importing diversity that mitigated the loss of strains through drift (Fig. S14, S15). Weakening NFDS did not have a substantial effect (Fig. S16, S17).

### Multi-locus NFDS and asymmetrical transformation shape species-wide phylogenies

Bacterial population structure is often analysed using trees constructed from core genome SNPs. As the distribution of these polymorphisms was affected by both transformation and multi-locus NFDS, neighbour-joining trees were inferred from the genomic data (Fig. S18) and simulation outputs (Fig. 6a), and compared using two statistics. The first, the phylogenetic diversity of the tree (the sum of branch lengths) per tip, represented the overall diversity of the population. The second, Pybus and Harvey’s γ, summarised tree shapes using the relative positioning of internal nodes [80]. Higher γ values correspond to internal branching events being closer to leaf nodes than expected under neutrality. The highest values were generated by the simulations without transformation, in which many clades were flat (Fig. 6b), as in the absence of diversification through recombination, identical genotypes were repeatedly sampled from the population.

**Figure 6:**
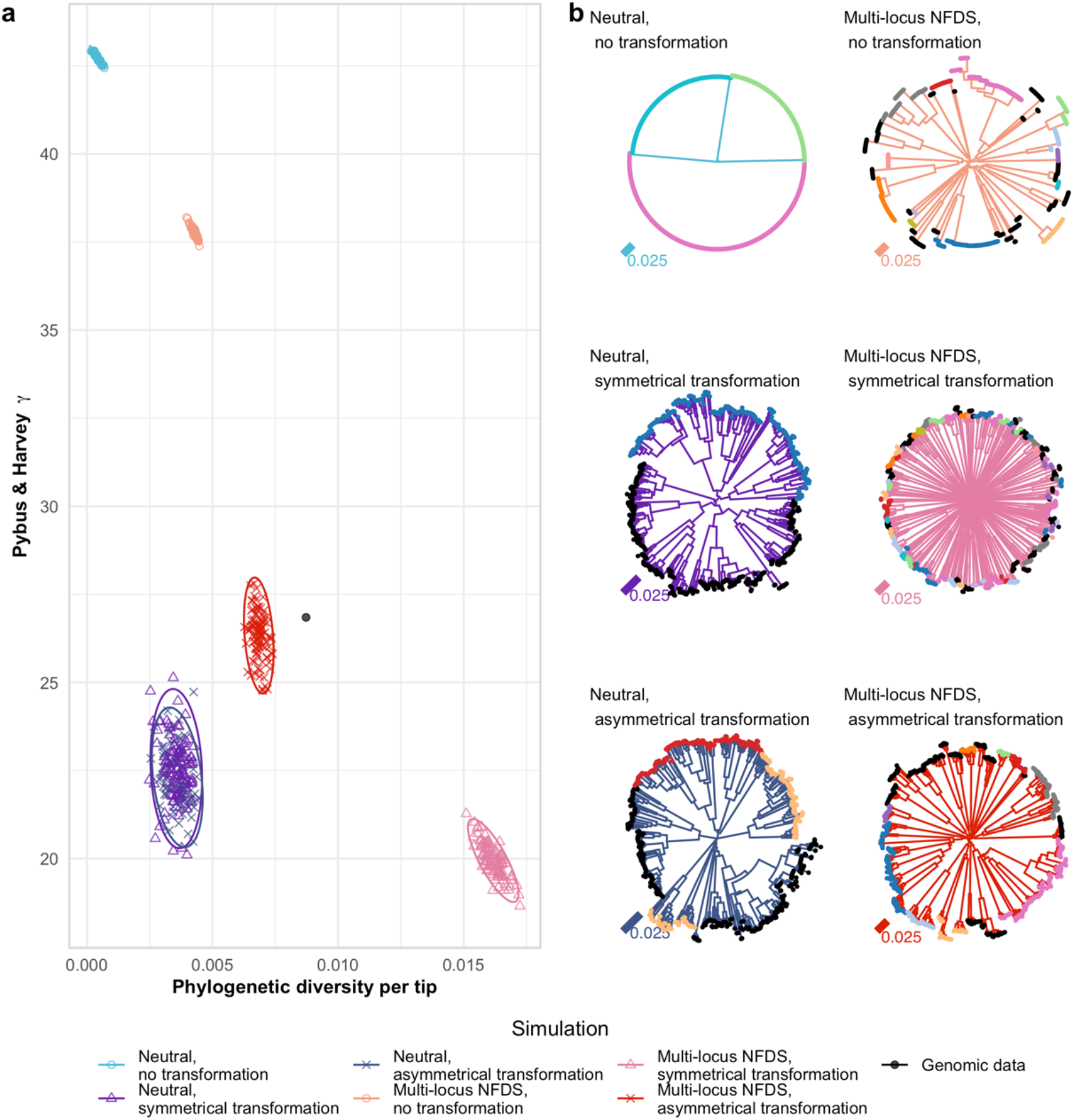
Comparison of trees between genomic data and simulation outputs. **a** Scatterplot comparing the characteristics of the neighbour-joining tree calculated from the core single nucleotide polymorphisms (*S* = 1,090) in the genomic data and those from the final timepoints of 100 replicate simulations. The horizontal axis quantifies the diversity of the population as phylogenetic diversity (sum of branch lengths) per tip (*N* = 616). The vertical axis quantifies the branching pattern as Pybus & Harvey’s γ. Point shapes and colours represent simulation types. Ellipses describe the distribution of each set of points assuming a multivariate t-distribution. **b** Representative trees corresponding to individual simulations from each parameter set. The branch colours indicate the simulation type, and the tip colours correspond to their original strain assignation in the genomic data. Strains that were rare in the genomic data were merged into a single category, which is marked by black tips.

The lowest γ values were associated with multi-locus NFDS simulations in which transformation was symmetrical. This reflects the high pairwise SNP distances generating star-like trees, with little internal structure. The many long branches mean these trees also have the highest phylogenetic diversity per tip. The trees describing neutrally-evolving populations exchanging sequence through transformation have similarly low γ values, but exhibited less phylogenetic diversity, reflecting the convergence of genotypes through recombination (Fig. 3). The trees most similar to that generated from the genomic data were generated by multi-locus NFDS simulations in which transformation was asymmetrical. These featured discernible clades separated by deep branches, previously regarded as characteristic of NFDS on a limited diversity of genotypes [88,89].

The tree statistics were sensitive to simulation conditions. Saltational transformation resulted in higher γ values, particularly for trees generated from multi-locus NFDS simulations featuring asymmetrical transformation, as flat clades of identical genotypes were more common (Fig. S19). Migration had little effect (Fig. S20). Weakening NFDS reduced the phylogenetic diversity per tip, and simulations in which selection did not prevent erosion of the accessory genome by asymmetrical transformation also failed to closely replicate the observed tree structure (Fig. S21).

## Discussion

Despite the extensive genotypic and phenotypic variation observed across diverse species such as *S. pneumoniae*, bacterial population genetic models have struggled to reject neutral models [27]. This reflects both the ability of sophisticated neutral models to reproduce particular aspects of bacterial populations [19,86], and the insensitivity of some statistics in assessing model fits [85,90]. Population genomics datasets enable multiple tests of model fitting, illustrated by the locus frequencies, genome sizes, pairwise distances, strain frequency and phylogenetic statistics employed in this study. These demonstrate the best-fitting models are multi-locus NFDS simulations with transformation absent or asymmetrical. In the absence of recombination, multi-locus NFDS stably preserves the original population (Fig. S22), without driving the oscillations possible with strong NFDS on few loci, as in models motivated by the dynamics of influenza [55,91]. However, the lack of transformation is neither consistent with the distribution of diversity in the starting population, nor the detection of interstrain recombinations [23,24]. When asymmetrical transformation is combined with multi-locus NFDS, the populations’ characteristics were preserved while permitting the genesis of new strains, divergent from the starting population (Fig. S22). This parameterisation is a more comprehensive explanation for the stable accessory locus and SNP frequencies, range of genome sizes, distribution of pairwise distances and tree structure in the observed data.

Asymmetric transformation and multi-locus NFDS interact to generate outbreeding depression that is not shared by other model parameterisations. The NFDS aspect selects for a diverse population through maintaining all intermediate-frequency accessory loci at their equilibrium frequencies. This is mediated by competition between genotypes with overlapping accessory locus profiles. When transformation is symmetrical, the majority of isolates are separated by pairwise distances similar in magnitude to those separating strains in the genomic data. This likely represents the simulation reaching an equilibrium at which accessory locus profiles least overlap, and therefore competition is minimised. Consistent with this hypothesis, the modal between-strain accessory distances in the genomic data are greater than those expected between random genotypes, under the assumption that the accessory genes have equilibrium frequencies (Fig. S23). Many NFDS models predict pathogens will exhibit a strong tendency towards continuous diversification [48]. However, asymmetrical transformation inhibits MSPs reaching this equilibrium, as when new genotypes are generated through sequence exchange, the recipient and recombinant are typically distinguished by the deletion of loci present in the recipient but absent in the donor. This reduces the instantaneous frequency of these loci below their equilibrium frequencies, thereby increasing the fitness of the unmodified recipient genotype (which still encodes these loci) relative to the recombinant progeny (from which these loci have been removed). As transformation generates novel individual genotypes, each so rare that they are liable to be lost through neutral genetic drift, even relatively weak selection can eliminate many such recombinants, and therefore preserve both accessory locus frequencies and MSP structures. Changes in SNP allele frequency are therefore indirectly limited both by selection against genotypes drifting, by multi-locus NFDS, and against transformation, through outbreeding depression eliminating recombinants. Yet the transformability of *S. pneumoniae* can nevertheless be advantageous overall, if it is asymmetric, because the cost of interstrain recombination can be outweighed by the benefits of ‘chromosomal curing’ through within-strain transformation [92].

The model makes many simplifications. No introduction of novel variation through mutation occurred, as only intermediate-frequency alleles were modelled. These are unlikely to have been recently generated, as they are shared between distant populations with divergent strain compositions [20]. Exchange of variation through transformation occurred at a single rate, rather than the range observed across the species [4,22], and was inferred from reconstructions of clinical isolates’ evolutionary history [70], which is necessarily measured after selection. Hence the actual pre-selection rate of transformation is higher, although this should not qualitatively alter the results, unless it is high enough to drive the elimination of some accessory loci. This rate was simulated as being uniform across loci and over time. This does not reflect the punctuated nature of interspecies recombination [70], which was approximated by simulations with ‘saltational’ transformation. This had little effect on simulations in which transformation was symmetrical. However, if it were asymmetrical, outbreeding depression was increased, as each recombinant deleted a greater number of loci, and consequently many genotypes never deviated from their initial form (Fig. S24). It could be that large, infrequent recombinations do not exhibit the same deletional bias as more common transformation events. Such a process would increase the rate of new strain formation, and therefore a mechanism driving outbreeding depression akin to that described here would be in greater need to preserve MSPs.

Many accessory loci are autonomously mobile [20], and therefore the net asymmetry of the recombination processes affecting their distribution will favour insertion over deletion. This could be simulated using a parameterisation of ϕ > 1. Assuming such elements were parasitic, recombinant progeny would likely be outcompeted by the original genotype, which does not harbour the element. This would be consistent with the long-standing hypothesis that the steady-state frequency of ‘selfish’ elements reflects a balance between their mobility and selection against infected genotypes [90].

Another simplification was that all loci evolved independently, without any consideration of physical linkage. Non-mobile accessory loci are clustered into genomic islands [12,20], which can be found in many combinations, consistent with the “modular selection” model of local epistasis [46]. The coherence of these distinct islands would decrease the number of possible accessory locus genotypes, and increase the variance and heterogeneity of the locus frequency, genome size and pairwise distance distributions generated by simulations featuring transformation. Additionally, as the extent of transformational asymmetry is determined by the size of a genomic island [68], and the selective pressure acting upon it will be determined by the functions of the encoded genes, this means the prevention of decay by NFDS will actually be a property of each island, rather than the population as a whole. Contrasting with the strong linkage between accessory loci within a genomic island, there is little species-wide linkage between these islands and the core genome SNPs that surround them in *S. pneumoniae* [20]. Hence there is limited evidence for localised “fronts” of diversification surrounding genomic islands, as was previously hypothesised to flank structural variation [37,87,94]. Therefore, the extent to which neutral SNP frequencies were preserved through selection on accessory loci should represent a minimum, that may be slightly higher in more realistic simulations that account for the limited linkage between some core and accessory variation.

Many of the population characteristics reproduced by the simulations featuring multi-locus NFDS and asymmetrical transformation are common to multiple species. Whether multi-locus NFDS can be applied to other microbes is unclear, but the functions encoded by the *S. pneumoniae* accessory genome hypothesised to be under NFDS are common across bacteria [20], and the model was consistent with the strain dynamics within an *Escherichia coli* population [95]. Although transformation is not ubiquitous across bacteria [92], the necessity for mechanisms blocking the integration of parasitic mobile elements is widespread [96,97]. For instance, restriction-modification systems have frequently been proposed to shape bacterial populations through blocking exchange of DNA [28,29,98], but recent experimental work has demonstrated they primarily inhibit the integration of novel genes, rather than the exchange of core genome variation [99,100]. This would impose a similar asymmetry on recombination at accessory loci as identified for transformation, suggesting multi-locus NFDS could still maintain MSPs through selecting against recombinants with reduced accessory genomes [28,29,98]. These processes are likely limited to accounting for diversity within species or subspecies, as the simulated genotypes compete within an homogeneous niche, and consequently the model does not account for ecological differentiation at higher taxonomic groupings [38,39]. Nevertheless, it is important to conclude that phenotypic variation between sets of bacterial genotypes should not be assumed to represent adaptation to distinct niches.

## Acknowledgements

G.L.H., J.A.L. and N.J.C. were supported by the UK Medical Research Council and Department for International Development (grant no. MR/R015600/1). N.J.C. was supported by a Sir Henry Dale Fellowship, jointly funded by Wellcome and the Royal Society (grant no. 104169/Z/14/A). J.C. was supported by European Research Council grant 742158. C.C. was supported by the Engineering and Physical Sciences Research Council of the United Kingdom (grant nos. EP/K026003/1 and EP/N014529/1) and the Government of Canada’s Canada 150 Research Chair programme. W.P.H. was supported by NIH grant R01 AI106786. M.L. was supported by the National Institute of General Medical Sciences (cooperative agreement U54GM088558). The content is solely the responsibility of the authors and does not necessarily represent the official views of the National Institute of General Medical Sciences or the NIH. Computational support was provided by the Imperial College Research Computing Service (DOI: 10.14469/hpc/2232).

## References

1. Russell JB. Discussion on Diphtheria. BMJ. 1891;2: 631–640.

2. Joseph FH. Notes on some Pathogenic Bacteria as found in the Transvaal, and the Variations from their European Prototype. Rep S African Assoc Adv Sci. 1904;2: 237–242.

3. Eyre JW, Washbourn JW. Varities and virulence of the pneumococcus. Lancet. 1899;153: 19–22. doi:10.1016/S0140-6736(01)78950-7

4. Gladstone RA, Lo SW, Lees JA, Croucher NJ, van Tonder AJ, Corander J, et al. International genomic definition of pneumococcal lineages, to contextualise disease, antibiotic resistance and vaccine impact. EBioMedicine. 2019;43: 338–346. doi:10.1016/j.ebiom.2019.04.021

5. Colijn C, Corander J, Croucher NJ. Designing ecologically optimized pneumococcal vaccines using population genomics. Nat Microbiol. 2020;5: 473–485. doi:10.1038/s41564-019-0651-y

6. Weinberger DM, Malley R, Lipsitch M. Serotype replacement in disease after pneumococcal vaccination. The Lancet. 2011. pp. 1962–1973. doi:10.1016/S0140-6736(10)62225-8

7. Lefevre JC, Faucon G, Sicard AM, Gasc AM. DNA fingerprinting of *Streptococcus pneumoniae* strains by pulsed-field gel electrophoresis. J Clin Microbiol. 1993;31: 2724 LP – 2728. Available: http://jcm.asm.org/content/31/10/2724.abstract

8. Selander RK, Levin BR. Genetic diversity and structure in *Escherichia coli* populations. Science. 1980;210: 545–547. doi:10.1126/science.6999623

9. Smith JM, Smith NH, O’Rourke M, Spratt BG. How clonal are bacteria? Proc Natl Acad Sci. 1993;90: 4384–4388.

10. Maiden MCJ, Bygraves JA, Feil E, Morelli G, Russell JE, Urwin R, et al. Multilocus sequence typing: A portable approach to the identification of clones within populations of pathogenic microorganisms. Proc Natl Acad Sci U S A. 1998;95: 3140–3145. doi:10.1073/pnas.95.6.3140

11. Smith JM, Feil EJ, Smith NH. Population structure and evolutionary dynamics of pathogenic bacteria. BioEssays. 2000;22: 1115–1122. doi:10.1002/1521-1878(200012)22:12<1115::AID-BIES9>3.0.CO;2-R

12. Croucher NJ, Coupland PG, Stevenson AE, Callendrello A, Bentley SD, Hanage WP. Diversification of bacterial genome content through distinct mechanisms over different timescales. Nat Commun. Nature Publishing Group; 2014;5: 5471.

13. Lees JA, Harris SR, Tonkin-Hill G, Gladstone RA, Lo S, Weiser JN, et al. Fast and flexible bacterial genomic epidemiology with PopPUNK. Genome Res. 2019;29: 304–316. doi:10.1101/360917

14. Alikhan NF, Zhou Z, Sergeant MJ, Achtman M. A genomic overview of the population structure of *Salmonella*. PLoS Genet. 2018;14: e1007261. doi:10.1371/journal.pgen.1007261

15. Hanage WP, Spratt BG, Turner KME, Fraser C. Modelling bacterial speciation. Philos Trans R Soc B Biol Sci. 2006;361: 2039–2044. doi:10.1098/rstb.2006.1926

16. Ford Doolittle W, Papke RT. Genomics and the bacterial species problem. Genome Biol. 2006;7: 116. doi:10.1186/gb-2006-7-9-116

17. Shapiro BJ, David LA, Friedman J, Alm EJ. Looking for Darwin’s footprints in the microbial world. Trends Microbiol. 2009;17: 196–204. doi:10.1016/j.tim.2009.02.002

18. Abu-Raddad LJ, Ferguson NM. The impact of cross-immunity, mutation and stochastic extinction on pathogen diversity. Proc R Soc B Biol Sci. 2004;271: 2431–2438. doi:10.1098/rspb.2004.2877

19. Fraser C, Hanage WP, Spratt BG. Neutral microepidemic evolution of bacterial pathogens. Proc Natl Acad Sci U S A. 2005;102: 1968–1973. doi:10.1073/pnas.0406993102

20. Corander J, Fraser C, Gutmann MU, Arnold B, Hanage WP, Bentley SD, et al. Frequency-dependent selection in vaccine-associated pneumococcal population dynamics. Nat Ecol Evol. 2017;1: 1950–1960. doi:10.1038/s41559-017-0337-x

21. Buckee CO, Jolley KA, Recker M, Penman B, Kriz P, Gupta S, et al. Role of selection in the emergence of lineages and the evolution of virulence in *Neisseria meningitidis*. Proc Natl Acad Sci U S A. 2008;105: 15082–15087. doi:10.1073/pnas.0712019105

22. Croucher NJ, Finkelstein JA, Pelton SI, Mitchell PK, Lee GM, Parkhill J, et al. Population genomics of post-vaccine changes in pneumococcal epidemiology. Nat Genet. 2013;45: 656–663. doi:10.1038/ng.2625

23. Croucher NJ, Hanage WP, Harris SR, McGee L, van der Linden M, de Lencastre H, et al. Variable recombination dynamics during the emergence, transmission and “disarming” of a multidrug-resistant pneumococcal clone. BMC Biol. 2014;12: 49. doi:10.1186/1741-7007-12-49

24. Croucher NJ, Chewapreecha C, Hanage WP, Harris SR, McGee L, van der Linden M, et al. Evidence for soft selective sweeps in the evolution of pneumococcal multidrug resistance and vaccine escape. Genome Biol Evol. Oxford University Press; 2014;6: 1589–1602. doi:10.1093/gbe/evu120

25. Vos M. A species concept for bacteria based on adaptive divergence. Trends Microbiol. 2011;19: 1–7. doi:10.1016/j.tim.2010.10.003

26. Whitaker RJ. Allopatric origins of microbial species. Philos Trans R Soc B Biol Sci. 2006;361: 1975–1984. doi:10.1098/rstb.2006.1927

27. Fraser C, Alm EJ, Polz MF, Spratt BG, Hanage WP. The bacterial species challenge: Making sense of genetic and ecological diversity. Science. 2009;323: 741–746. doi:10.1126/science.1159388

28. Hanage WP, Fraser C, Spratt BG. Fuzzy species among recombinogenic bacteria. BMC Biol. 2005;3: 6. doi:10.1186/1741-7007-3-6

29. Fraser C, Hanage WP, Spratt BG. Recombination and the nature of bacterial speciation. Science. 2007;315: 476–480. doi:10.1126/science.1127573

30. Sheppard SK, McCarthy ND, Falush D, Maiden MCJ. Convergence of Campylobacter species: Implications for bacterial evolution. Science. 2008;320: 237–239. doi:10.1126/science.1155532

31. Laible G, Spratt BG, Hakenbeck R. Interspecies recombinational events during the evolution of altered PBP 2x genes in penicillin-resistant clinical isolates of *Streptococcus pneumoniae*. Mol Microbiol. 1991;5: 1993–2002. doi:10.1111/j.1365-2958.1991.tb00821.x

32. Dowson CG, Coffey TJ, Kell C, Whiley RA. Evolution of penicillin resistance in *Streptococcus pneumoniae;* the role of *Streptococcus mitis* in the formation of a low affinity PBP2B in *S. pneumoniae*. Mol Microbiol. 1993/08/01. 1993;9: 635–643. doi:10.1111/j.1365-2958.1993.tb01723.x

33. Majewski J, Zawadzki P, Pickerill P, Cohan FM, Dowson CG. Barriers to genetic exchange between bacterial species: *Streptococcus pneumoniae* transformation. J Bacteriol. 2000;182: 1016–1023. doi:10.1128/JB.182.4.1016-1023.2000

34. Bobay L-M, Ochman H. Biological Species Are Universal across Life’s Domains. Genome Biol Evol. 2017;9: 491–501. doi:10.1093/gbe/evx026

35. Croucher NJ, Harris SR, Barquist L, Parkhill J, Bentley SD. A high-resolution view of genome-wide pneumococcal transformation. PLoS Pathog. 2012;8: e1002745. doi:10.1371/journal.ppat.1002745

36. Cohan F. Bacterial species and speciation. Syst Biol. 2001;50: 513–524. doi:10.1080/10635150118398

37. Nosil P, Funk DJ, Ortiz-Barrientos D. Divergent selection and heterogeneous genomic divergence. Mol Ecol. 2009;18: 375–402. doi:10.1111/j.1365-294X.2008.03946.x

38. Cohan FM, Perry EB. A Systematics for Discovering the Fundamental Units of Bacterial Diversity. Curr Biol. 2007;17: R373–R386. doi:10.1016/j.cub.2007.03.032

39. Marttinen P, Hanage WP. Speciation trajectories in recombining bacterial species. PLoS Comput Biol. 2017;13: e1005640. doi:10.1371/journal.pcbi.1005640

40. Gjini E, Valente C, Sá-Leão R, Gomes MGM. How direct competition shapes coexistence and vaccine effects in multi-strain pathogen systems. J Theor Biol. 2016;388: 50–60. doi:10.1016/j.jtbi.2015.09.031

41. Numminen E, Cheng L, Gyllenberg M, Corander J. Estimating the Transmission Dynamics of *Streptococcus pneumoniae* from Strain Prevalence Data. Biometrics. 2013;69: 748–757. doi:10.1111/biom.12040

42. Majewski J, Cohan FM. Adapt globally, act locally: The effect of selective sweeps on bacterial sequence diversity. Genetics. 1999;152: 1459–1474.

43. Wiedenbeck J, Cohan FM. Origins of bacterial diversity through horizontal genetic transfer and adaptation to new ecological niches. FEMS Microbiol Rev. 2011;35: 957–976. doi:10.1111/j.1574-6976.2011.00292.x

44. Barraclough TG, Balbi KJ, Ellis RJ. Evolving Concepts of Bacterial Species. Evol Biol. 2012;39: 148–157. doi:10.1007/s11692-012-9181-8

45. Chesson P. Mechanisms of maintenance of species diversity. Annu Rev Ecol Syst. 2000;31: 343–366. doi:10.1146/annurev.ecolsys.31.1.343

46. Neher RA, Shraiman BI. Competition between recombination and epistasis can cause a transition from allele to genotype selection. Proc Natl Acad Sci U S A. 2009;106: 6866–6871. doi:10.1073/pnas.0812560106

47. Mostowy R, Croucher NJ, Andam CP, Corander J, Hanage WP, Marttinen P. Efficient Inference of Recent and Ancestral Recombination within Bacterial Populations. Mol Biol Evol. 2017;34: 1167–1182. doi:10.1093/molbev/msx066

48. Ferguson NM, Galvanl AP, Bush RM. Ecological and immunological determinants of influenza evolution. Nature. 2003;422: 428–433. doi:10.1038/nature01509

49. Cobey S, Lipsitch M. Niche and neutral effects of acquired immunity permit coexistence of pneumococcal serotypes. Science. 2012;335: 1376–1380.

50. Cobey S. Pathogen evolution and the immunological niche. Ann N Y Acad Sci. 2014;1320: 1–15. doi:10.1111/nyas.12493

51. Levin BR. Frequency-dependent selection in bacterial populations. Philos Trans R Soc Lond B Biol Sci. 1988;319: 459–72. doi:10.1098/rstb.1988.0059

52. Gomes MGM, Medley GF, Nokes DJ. On the determinants of population structure in antigenically diverse pathogens. Proc R Soc London Ser B Biol Sci. 2002;269: 227–233. doi:10.1098/rspb.2001.1869

53. Binsker U, Lees JA, Hammond AJ, Weiser JN. Immune exclusion by naturally acquired secretory IgA against pneumococcal pilus-1. J Clin Invest. 2020;130: 927–941. doi:10.1172/JCI132005

54. Gupta S, Maiden MCJ, Feavers IM, Nee S, May RM, Anderson RM. The maintenance of strain structure in populations of recombining infectious agents. Nat Med. 1996;2: 437–442.

55. Gupta S, Ferguson N, Anderson R. Chaos, persistence, and evolution of strain structure in antigenically diverse infectious agents. Science. 1998;280: 912–915.

56. Croucher NJ, Campo JJ, Le TQ, Liang X, Bentley SD, Hanage WP, et al. Diverse evolutionary patterns of pneumococcal antigens identified by pangenome-wide immunological screening. Proc Natl Acad Sci U S A. 2017;114: E357–E366. doi:10.1073/pnas.1613937114

57. Weinberger DM, Dagan R, Givon-Lavi N, Regev-Yochay G, Malley R, Lipsitch M. Epidemiologic Evidence for Serotype-Specific Acquired Immunity to Pneumococcal Carriage. J Infect Dis. 2008;197: 1511–1518. doi:10.1086/587941

58. Malley R, Lipsitch M, Bogaert D, Thompson CM, Hermans P, Claire Watkins A, et al. Serum Antipneumococcal Antibodies and Pneumococcal Colonization in Adults with Chronic Obstructive Pulmonary Disease. J Infect Dis. 2007;196: 928–935. doi:10.1086/520937

59. Campo JJ, Le TQ, Pablo J V, Hung C, Teng AA, Tettelin H, et al. Panproteome-wide analysis of antibody responses to whole cell pneumococcal vaccination. Elife. eLife Sciences Publications Limited; 2018;7: e37015. doi:10.7554/elife.37015

60. Buckee CO, Koelle K, Mustard MJ, Gupta S. The effects of host contact network structure on pathogen diversity and strain structure. Proc Natl Acad Sci U S A. 2004;101: 10839 LP – 10844. doi:10.1073/pnas.0402000101

61. Buckee CO, Recker M, Watkins ER, Gupta S. Role of stochastic processes in maintaining discrete strain structure in antigenically diverse pathogen populations. Proc Natl Acad Sci. 2011; 108: 15504 LP – 15509. doi:10.1073/pnas.1102445108

62. Watkins ER, Penman BS, Lourenço J, Buckee CO, Martin C, Maiden MCJ, et al. Vaccination Drives Changes in Metabolic and Virulence Profiles of *Streptococcus pneumoniae*. PLoS Pathog. 2015;11: e1005034. doi:10.1371/journal.ppat.1005034

63. Skwark MJ, Croucher NJ, Puranen S, Chewapreecha C, Pesonen M, Xu YY, et al. Interacting networks of resistance, virulence and core machinery genes identified by genome-wide epistasis analysis. PLoS Genet. 2017;13: e1006508. doi:10.1371/journal.pgen.1006508

64. Pensar J, Puranen S, Arnold B, MacAlasdair N, Kuronen J, Tonkin-Hill G, et al. Genome-wide epistasis and co-selection study using mutual information. Nucleic Acids Res. 2019;47: e112. doi:10.1093/nar/gkz656

65. Rocha EPC. Neutral Theory, Microbial Practice: Challenges in Bacterial Population Genetics. Mol Biol Evol. 2018;35: 1338–1347. doi:10.1093/molbev/msy078

66. Azarian T, Martinez PPP, Arnold BJ, Grant LR, Corander J, Fraser C, et al. Prediction of post-vaccine population structure of *Streptococcus pneumoniae* using accessory gene frequencies. bioRxiv. 2018; doi.org/10.1101/420315.

67. Johnston C, Campo N, Bergé MJ, Polard P, Claverys JP. *Streptococcus pneumoniae*, le transformiste. Trends Microbiol. 2014;22: 113–9. doi:10.1016/j.tim.2014.01.002

68. Apagyi KJ, Fraser C, Croucher NJ. Transformation asymmetry and the evolution of the bacterial accessory genome. Mol Biol Evol. 2018;35: 575–581. doi:10.1093/gbe/evs009

69. Croucher NJ, Walker D, Romero P, Lennard N, Paterson GK, Bason NC, et al. Role of conjugative elements in the evolution of the multidrug-resistant pandemic clone *Streptococcus pneumoniae^Spain23F^* ST81. J Bacteriol. 2008/12/31. 2009; 191: 1480–1489. doi:10.1128/jb.01343-08

70. Mostowy R, Croucher NJ, Hanage WP, Harris SR, Bentley S, Fraser C. Heterogeneity in the Frequency and Characteristics of Homologous Recombination in Pneumococcal Evolution. PLoS Genet. 2014;10: e1004300. doi:10.1371/journal.pgen.1004300

71. Croucher NJ, Klugman KP. The emergence of bacterial “Hopeful monsters.” MBio. 2014;5. doi:10.1128/mBio.01550-14

72. Kareiva P. Population dynamics in spatially complex environments: theory and data. Philos Trans - R Soc London, B. 1990;330: 175–190. doi:10.1098/rstb.1990.0191

73. R Core Team. R: A Language and Environment for Statistical Computing. R Found Stat Comput. Vienna: R Foundation for Statistical Computing; 2019; doi:10.1007/978-3-540-74686-7

74. Wickham H, Averick M, Bryan J, Chang W, McGowan L, François R, et al. Welcome to the Tidyverse. J Open Source Softw. 2019;4: 1686. doi:10.21105/joss.01686

75. Kassambara A. ggpubr: “ggplot2” Based Publication Ready Plots [Internet]. 2020. Available: https://cran.r-project.org/package=ggpubr

76. Blaser N. rdist: Calculate Pairwise Distances [Internet]. 2018. Available: https://cran.r-project.org/package=rdist

77. Spiess A-N. propagate: Propagation of Uncertainty [Internet]. 2018. Available: https://cran.r-project.org/package=propagate

78. Paradis E, Schliep K. ape 5.0: an environment for modern phylogenetics and evolutionary analyses in R. Bioinformatics. 2018;35: 526–528.

79. Yu G, Smith DK, Zhu H, Guan Y, Lam TTY. ggtree: an R package for visualization and annotation of phylogenetic trees with their covariates and other associated data. Methods Ecol Evol. 2017;8: 28–36. doi:10.1111/2041-210X.12628

80. Pybus OG, Harvey PH. Testing macro-evolutionary models using incomplete molecular phylogenies. Proc R Soc B Biol Sci. 2000;267: 2267–2272. doi:10.1098/rspb.2000.1278

81. Revell LJ. phytools: An R package for phylogenetic comparative biology (and other things). Methods Ecol Evol. 2012;3: 217–223. doi:10.1111/j.2041-210X.2011.00169.x

82. Csardi G, Nepusz T. The igraph software package for complex network research. InterJournal. 2006;Complex Sy.

83. Dixon P. VEGAN, a package of R functions for community ecology. J Veg Sci. 2003;14: 927–930. doi:10.1111/j.1654-1103.2003.tb02228.x

84. Kimura M. On the probability of fixation of mutant genes in a population. Genetics. 1962;47: 713–719.

85. Hanage WP, Fraser C, Spratt BG. The impact of homologous recombination on the generation of diversity in bacteria. J Theor Biol. 2006;239: 210–219. doi:10.1016/j.jtbi.2005.08.035

86. Marttinen P, Gutmann MU, Croucher NJ, Corander J, Hanage WP. Recombination produces coherent bacterial species clusters in both core and accessory genomes. Microb Genomics. 2015;1: e000038. doi:10.1099/mgen.0.000038

87. Lawrence JG. Gene Transfer in Bacteria: Speciation without Species? Theor Popul Biol. 2002;61: 449–460. doi:10.1006/tpbi.2002.1587

88. Barraclough TG, Birky CW, Burt A. Diversification in sexual and asexual organisms. Evolution (N Y). 2003;57: 2166–2172. doi:10.1111/j.0014-3820.2003.tb00394.x

89. Hudson RR. Gene genealogies and the coalescent process. Oxford Surveys in Evolutionary Biology. 1990. pp. 1–44.

90. Numminen E, Gutmann M, Shubin M, Marttinen P, Méric G, van Schaik W, et al. The impact of host metapopulation structure on the population genetics of colonizing bacteria. J Theor Biol. 2016;396: 53–62. doi:https://doi.org/10.1016/j.jtbi.2016.02.019

91. Castillo-Chavez C, Hethcote HW, Andreasen V, Levin SA, Liu WM. Epidemiological models with age structure, proportionate mixing, and cross-immunity. J Math Biol. 1989;27: 233–258. doi:10.1007/BF00275810

92. Croucher NJ, Mostowy R, Wymant C, Turner P, Bentley SD, Fraser C. Horizontal DNA Transfer Mechanisms of Bacteria as Weapons of Intragenomic Conflict. Barton NH, editor. PLOS Biol. 2016;14: e1002394. doi:10.1371/journal.pbio.1002394

93. Orgel LE, Crick FHC. Selfish DNA: The ultimate parasite. Nature. 1980;284: 604–7. doi:10.1038/284604a0

94. Vetsigian K, Goldenfeld N. Global divergence of microbial genome sequences mediated by propagating fronts. Proc Natl Acad Sci U S A. 2005;102: 7332–7337. doi:10.1073/pnas.0502757102

95. McNally A, Kallonen T, Connor C, Abudahab K, Aanensen D, Horner C, et al. Signatures of negative frequency dependent selection in colonisation factors and the evolution of a multi-drug resistant lineage of *Escherichia coli*. MBio. 2019;10: e00644–19. Available: http://biorxiv.org/content/early/2018/08/25/400374.abstract

96. Kwun MJ, Oggioni MR, Bentley SD, Fraser C, Croucher NJ. Synergistic Activity of Mobile Genetic Element Defences in *Streptococcus pneumoniae*. Genes (Basel). 2019;10: 707.

97. Iranzo J, Puigbo P, Lobkovsky AE, Wolf YI, Koonin E V. Inevitability of genetic parasites. Genome Biol Evol. 2016; doi:10.1093/gbe/evw193

98. Budroni S, Siena E, Dunning Hotopp JC, Seib KL, Serruto D, Nofroni C, et al. *Neisseria meningitidis* is structured in clades associated with restriction modification systems that modulate homologous recombination. Proc Natl Acad Sci U S A. 2011;108: 4494–4499. doi:10.1073/pnas.1019751108

99. Johnston C, Martin B, Granadel C, Polard P, Claverys JP. Programmed Protection of Foreign DNA from Restriction Allows Pathogenicity Island Exchange during Pneumococcal Transformation. PLoS Pathog. 2013;9: e1003178. doi:10.1371/journal.ppat.1003178

100. Kwun MJ, Oggioni MR, De Ste Croix M, Bentley SD, Croucher NJ. Excision-reintegration at a pneumococcal phase-variable restriction-modification locus drives within- and between-strain epigenetic differentiation and inhibits gene acquisition. Nucleic Acids Res. 2018;46: 11438–11453. doi:10.1093/nar/gky906

