## Supplementary Figures for "Negative frequency-dependent selection and asymmetrical transformation stabilise multi-strain bacterial population structures"

**Supplementary Information**

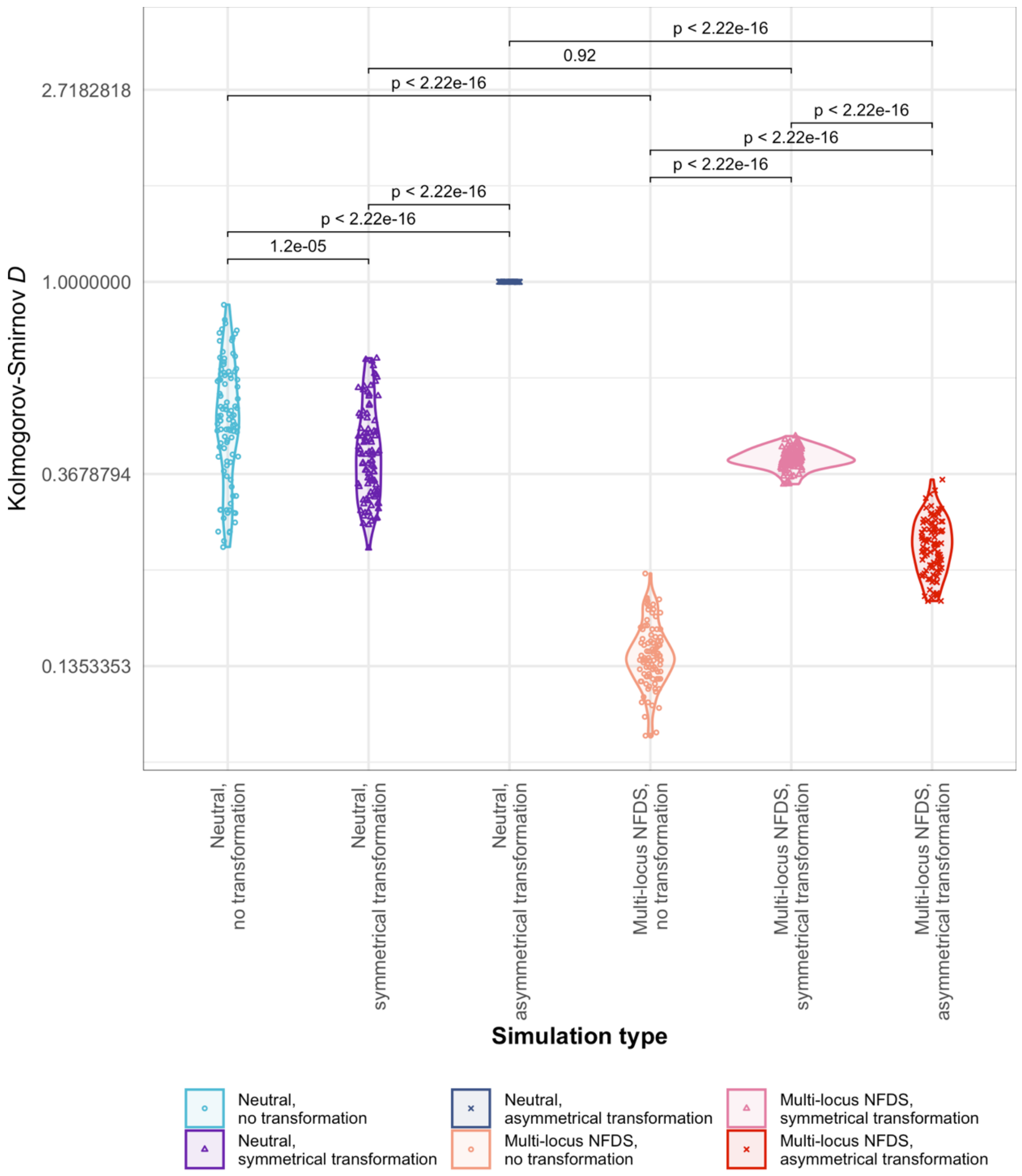


**Figure S1**: Violin plots comparing the observed distribution of intermediate-frequency loci per genome to those from the final timestep of simulations without migration (Fig. 2). The deviation between the observed and simulated data ($N$ = 100 replicates for each parameter combination) was measured using the Kolmogorov-Smirnov $D$ statistic. The values for each individual simulation are shown by points, and summarised over replicates as a violin plot. Pairwise Wilcoxon rank-sum tests were then undertaken between the $D$ statistics from sets of simulations that differed only in the mode of transformation or selection. The $p$ values from these tests are annotated at the top of the chart.

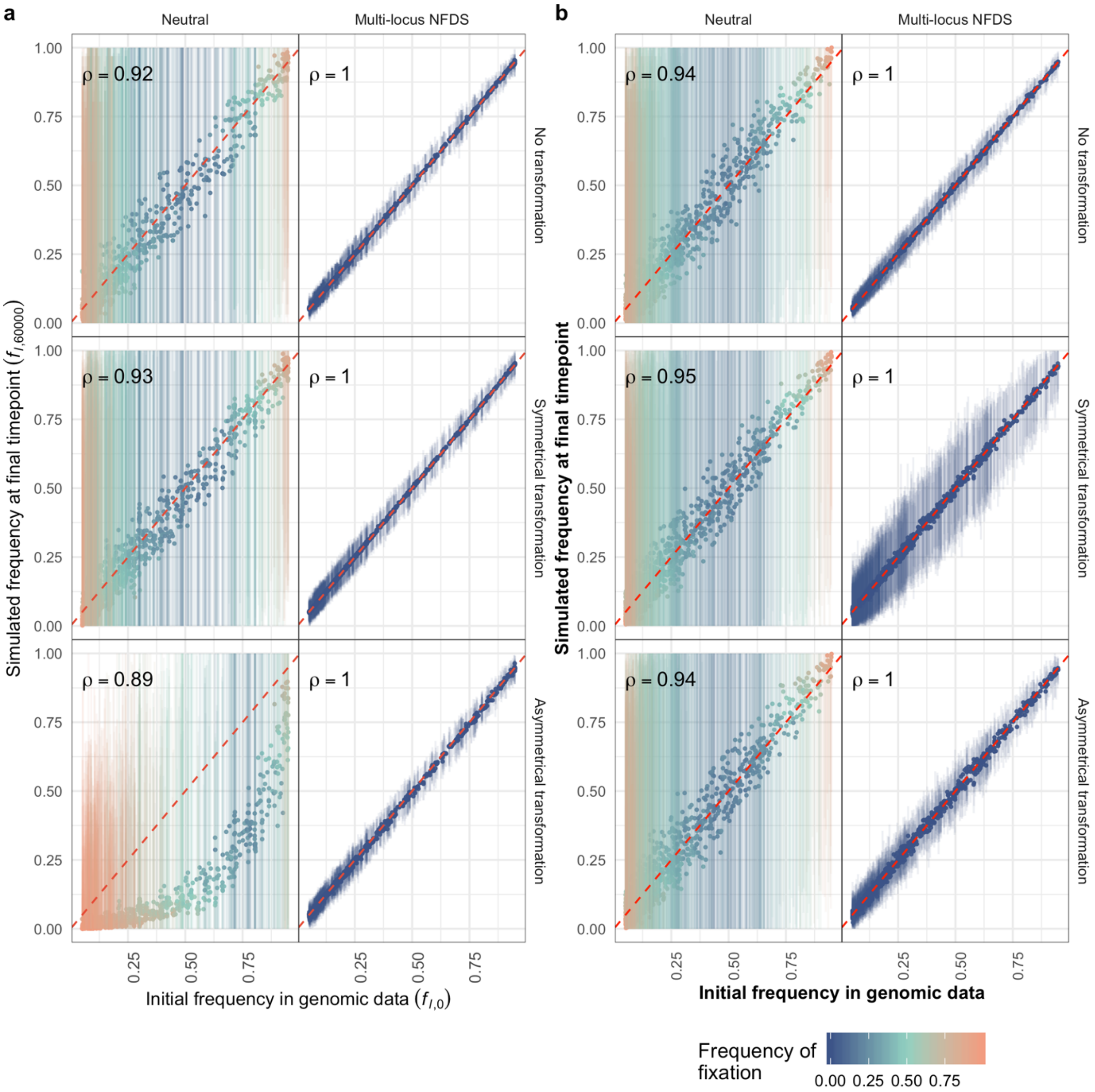


**Figure S2**: Scatterplots comparing the frequency of alleles in the initial timepoint in the genomic data to their frequency in the final simulation timepoint ($N$ = 616 isolates sampled from each simulation). Data are displayed as in Fig. 1. These simulations featured saltational transformation (Table 1).

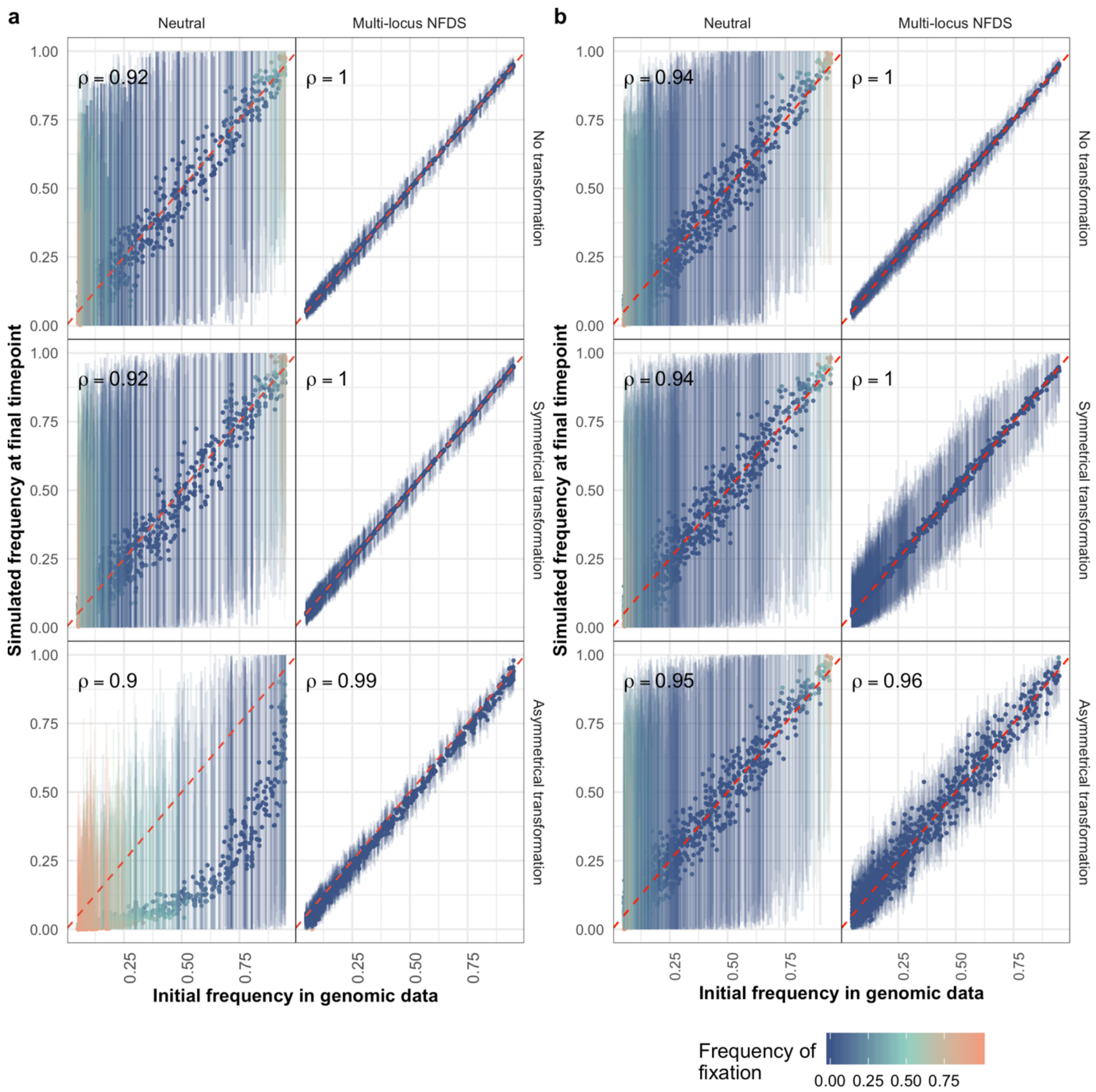


**Figure S3**: Scatterplots comparing the frequency of alleles at the initial timepoint in the genomic data to their frequency in the final simulation timepoint ($N$ = 616 isolates sampled from each simulation). Data are displayed as in Fig. 1. These simulations featured inward migration (Table 1).

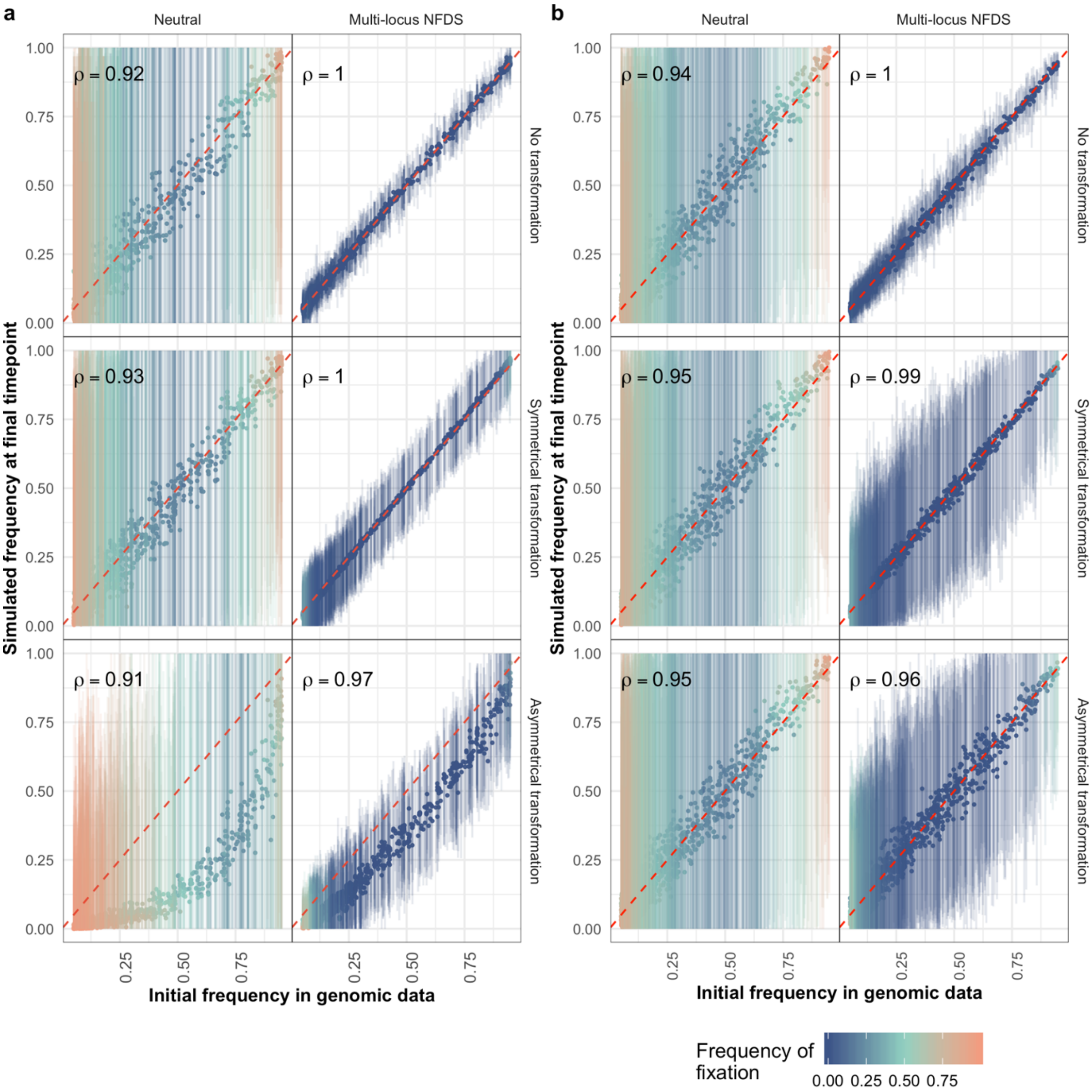


**Figure S4**: Scatterplots comparing the frequency of alleles at the initial timepoint in the genomic data to their frequency in the final simulation timepoint (*N* = 616 isolates sampled from each simulation). Data are displayed as in Fig. 1. These simulations featured weak multi-locus NFDS (Table 1).

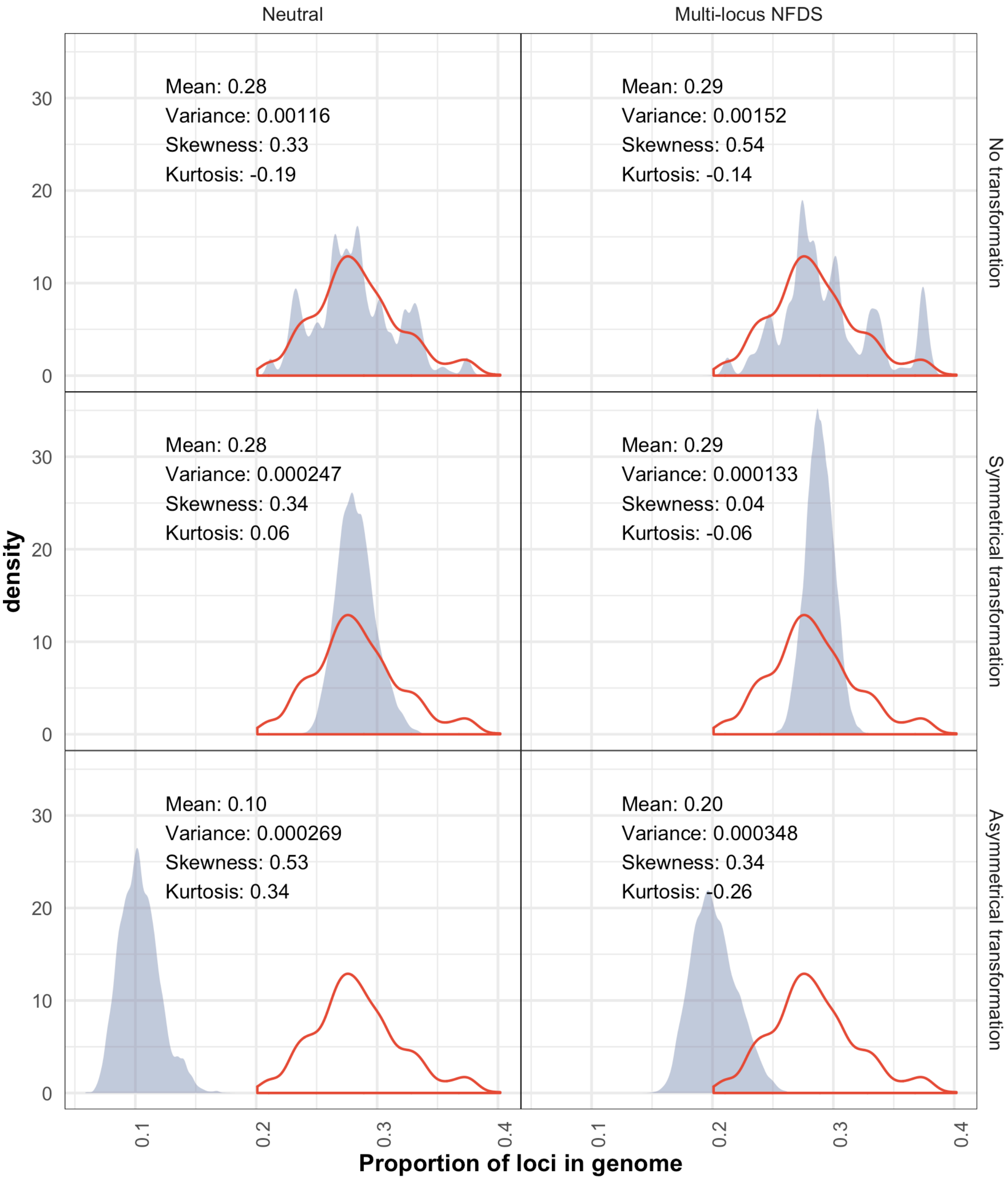


**Figure S5:** Density plots comparing the distribution of the number of accessory loci per isolate in the genomic data with those from the final timepoint of simulations. Data are displayed as in Fig. 2. These simulations featured weak multi-locus NFDS (Table 1).

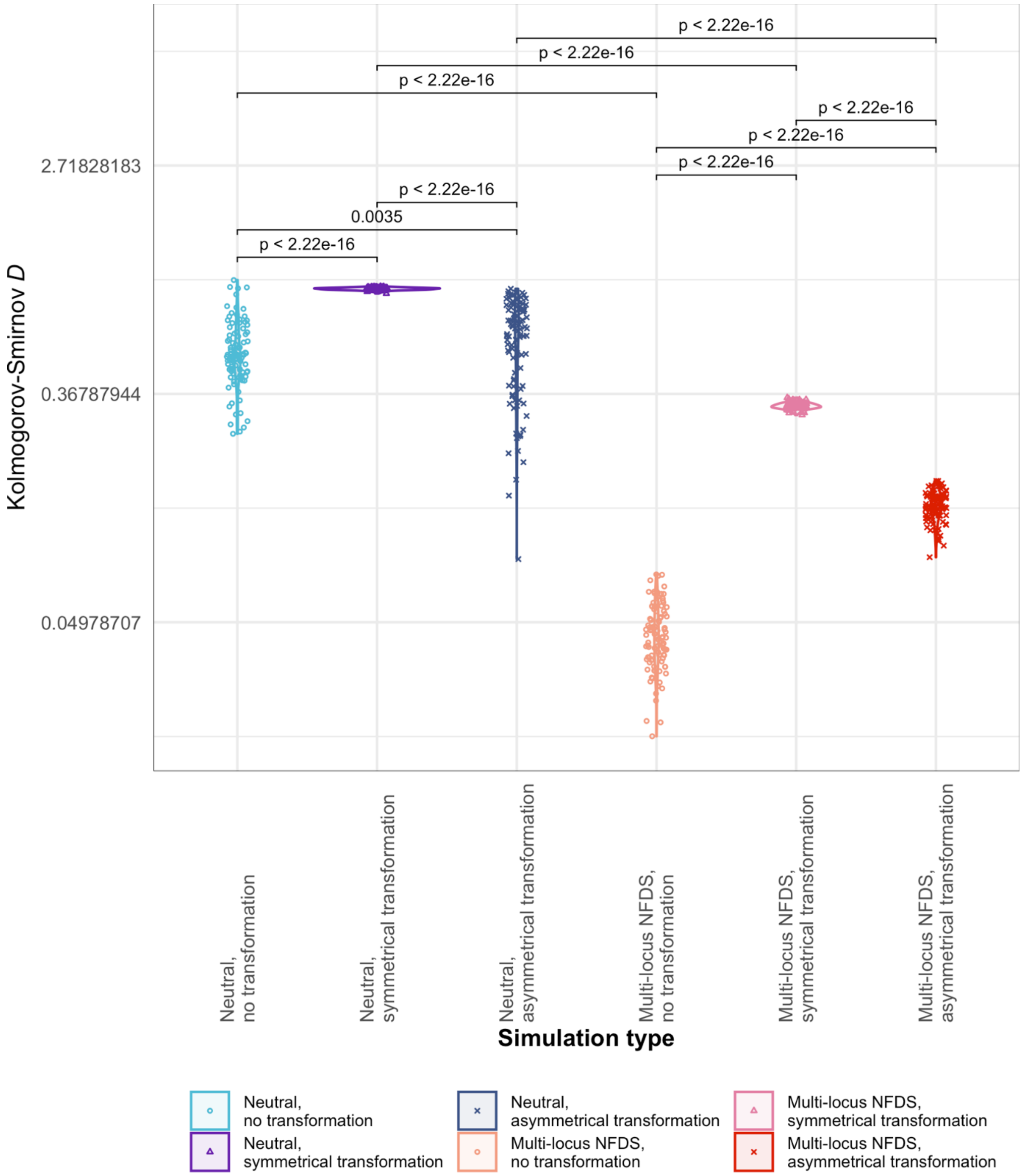


**Figure S6**: Violin plots comparing the observed distribution of pairwise Jaccard distances, calculated from the accessory loci encoded by genomes, with those from the final timestep of simulations without migration (Fig. 3). Data are shown as in Fig. S1.

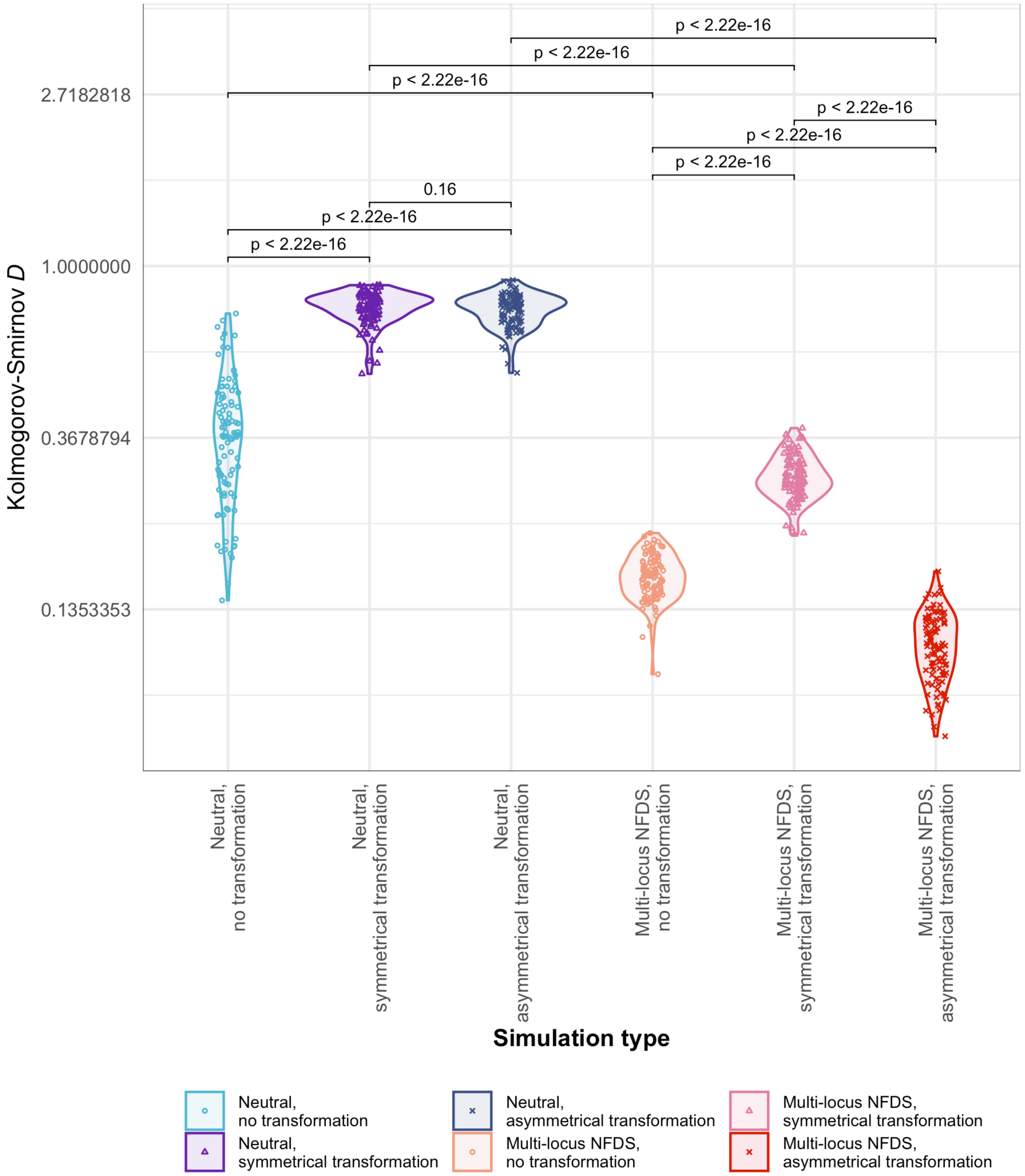


**Figure S7**: Violin plots comparing the observed distribution of pairwise Hamming distances, calculated from core genome SNPs, to those from the final timesteps of simulations without migration (Fig. 3). Data are shown as in Fig. S1.

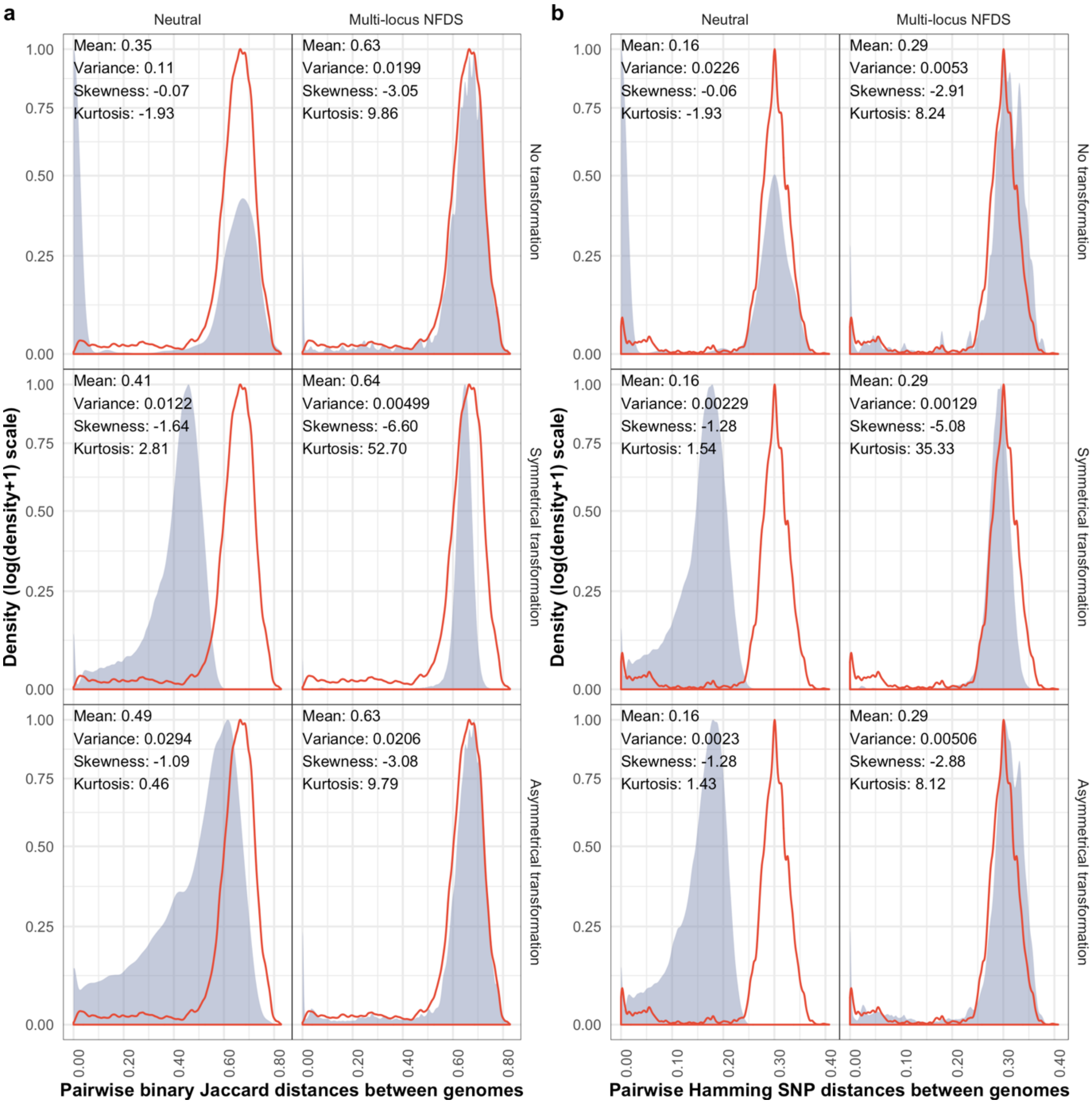


**Figure S8:** Density plots comparing the distributions of pairwise genetic distances between isolates in the genome data and at the final timepoint of simulations. Data are displayed as in Fig. 3. These simulations featured saltational transformation (Table 1).

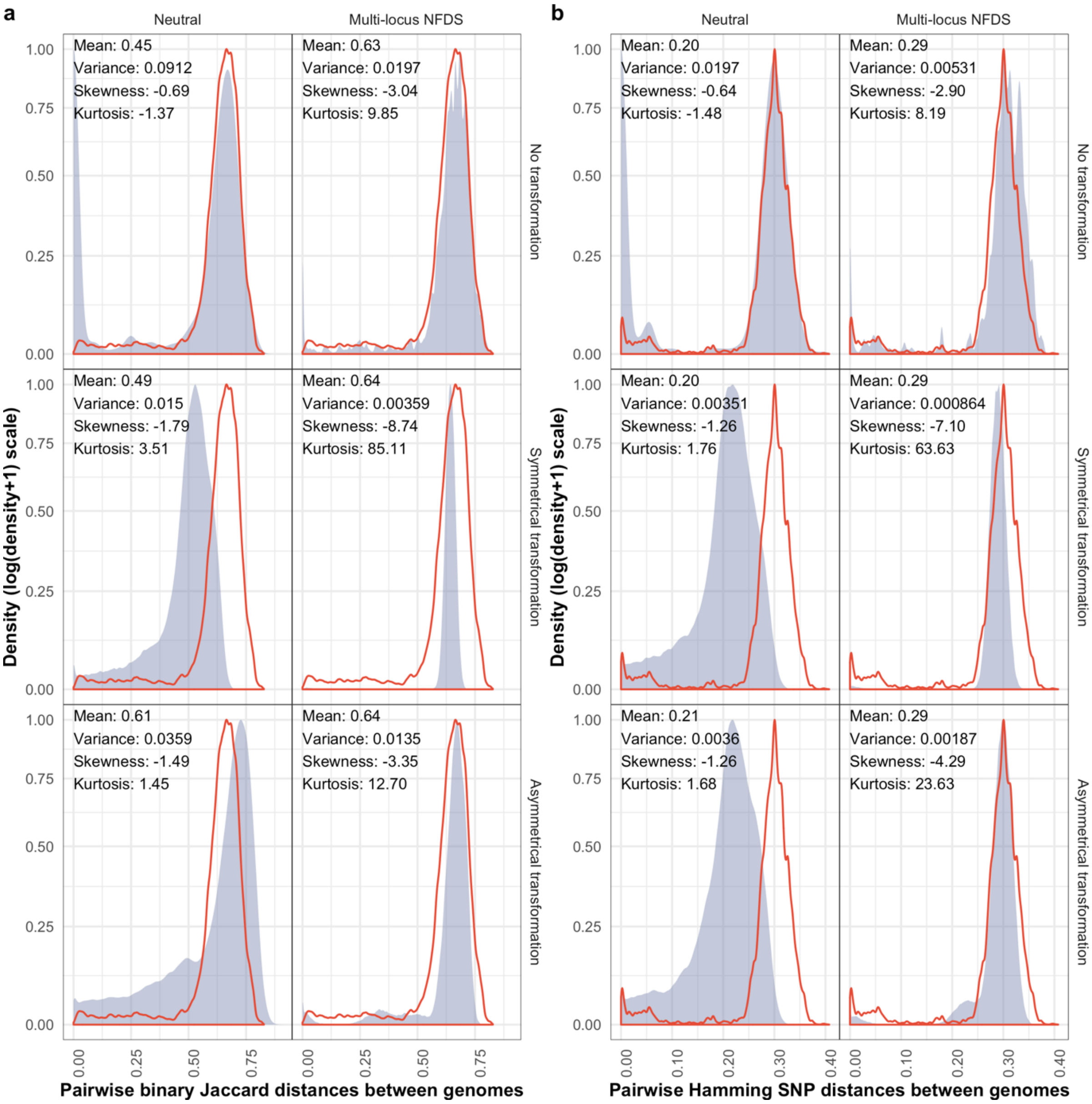


**Figure S9:** Density plots comparing the distributions of pairwise genetic distances between isolates in the genome data and at the final timepoint of simulations. Data are displayed as in Fig. 3. These simulations featured inward migration (Table 1).

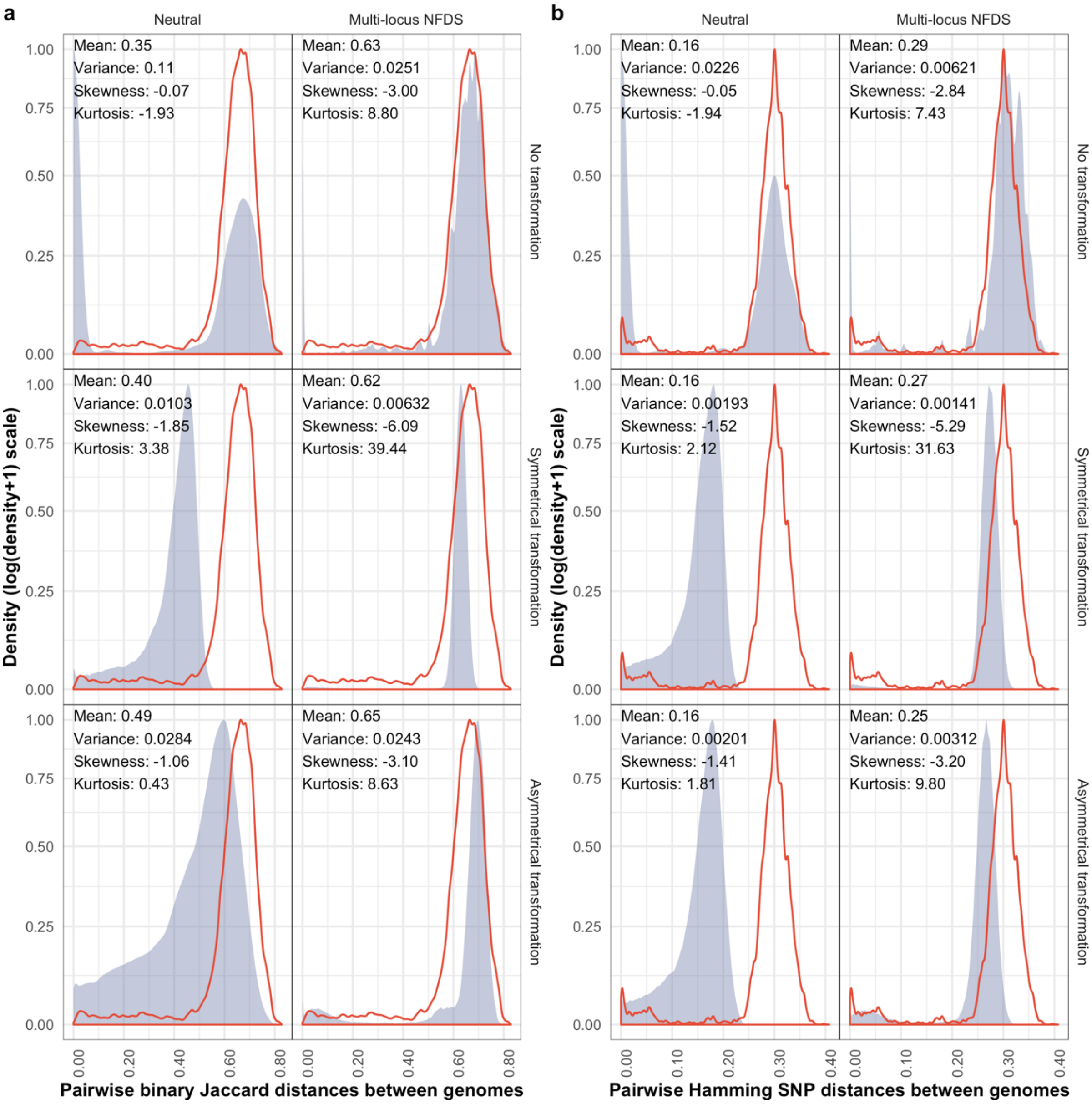


**Figure S10:** Density plots comparing the distributions of pairwise genetic distances between isolates in the genome data and at the final timepoint of simulations. Data are displayed as in Fig. 3. These simulations featured weak multi-locus NFDS (Table 1).

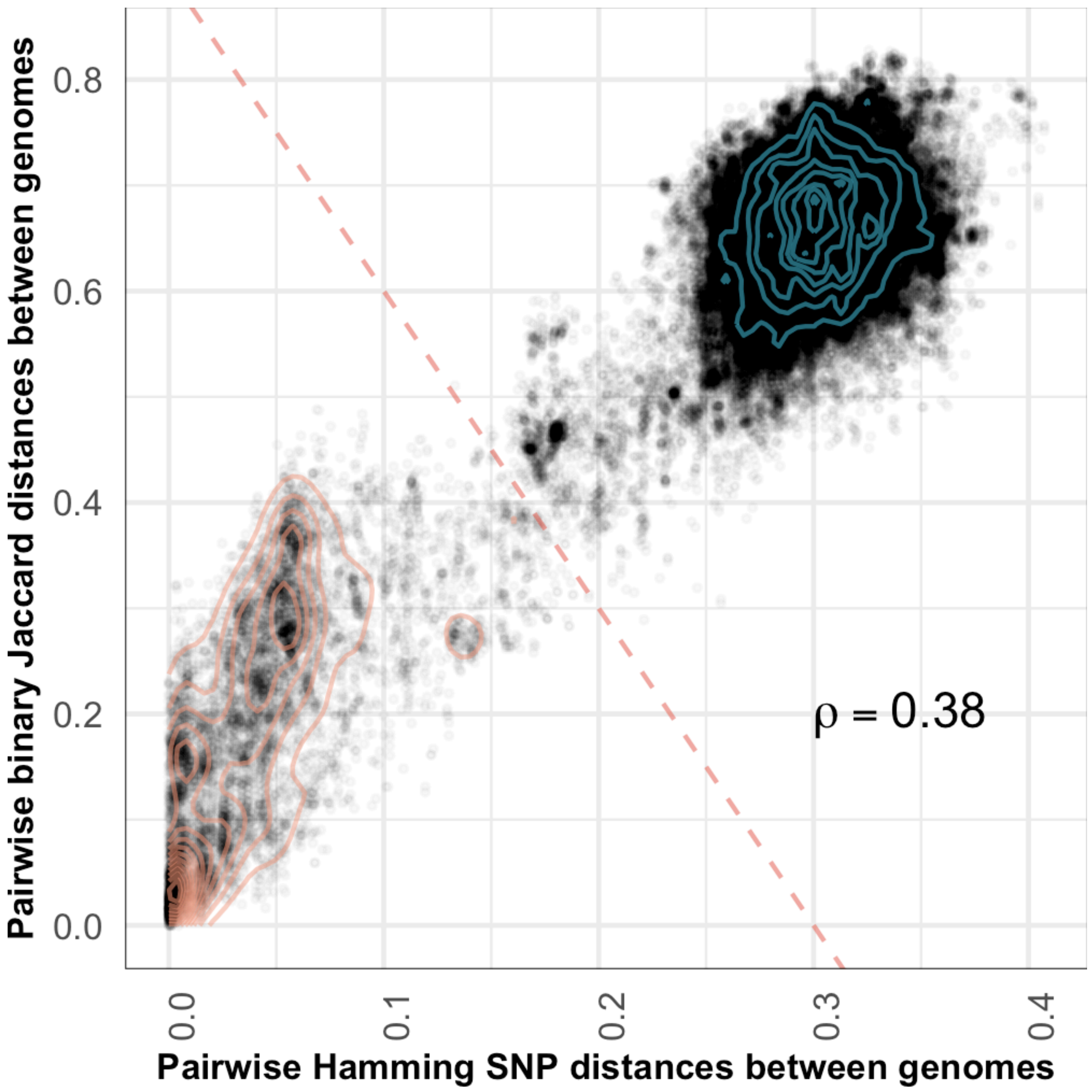


**Figure S11:** Scatterplot showing the genetic distances (*N* = 189,420) between isolates in the genomic data, with the horizontal axis representing divergence in core genome single nucleotide polymorphisms (*S* = 1090), and the vertical axis representing divergence in accessory loci (*L* = 1090). The red diagonal is a threshold distinguishing within- and between-strain distances. The contours summarise the distribution of within- and between-strain pairwise distances shown by the points (orange and blue lines, respectively).

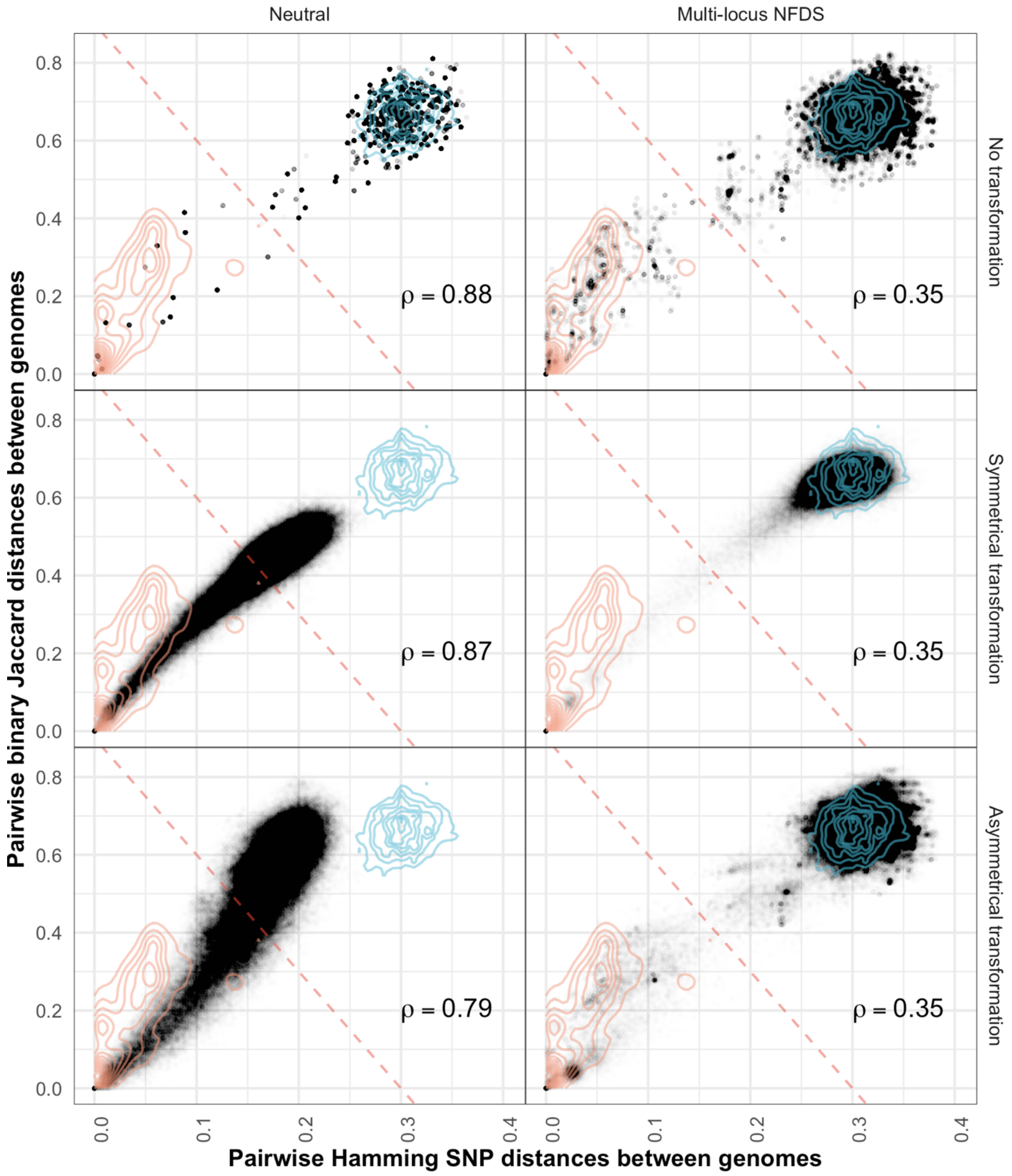


**Figure S12:** Scatterplots comparing the distributions of pairwise genetic distances between isolates at the final timepoint of simulations. Data are shown as in Fig. 4. These simulations featured saltational transformation (Table 1).

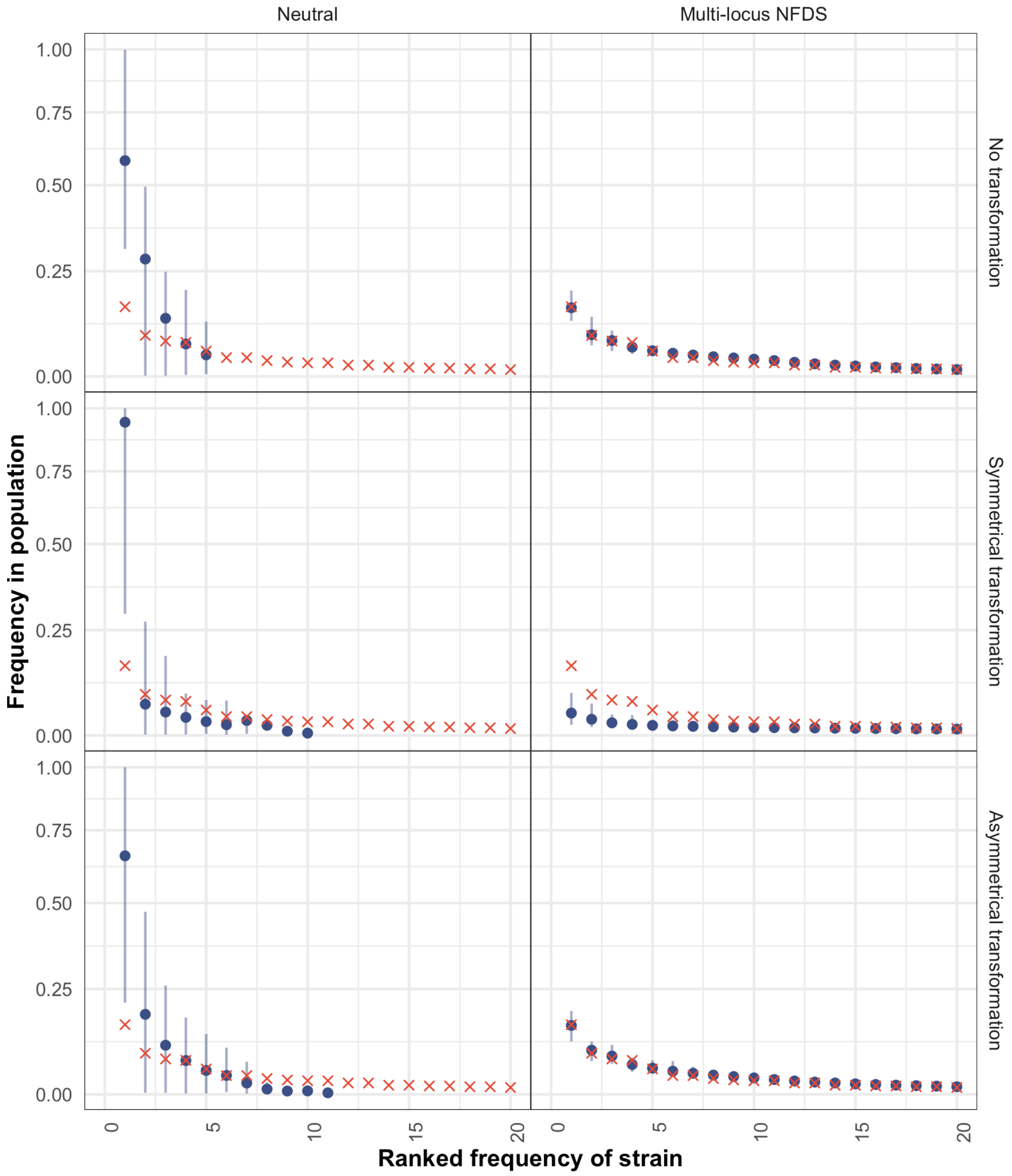


**Figure S13:** Scatterplots comparing the rank-frequency distributions of strains in the overall set of genomic data (red crosses) and those from samples of isolates from the final timepoint of 100 replicate simulations (blue points). Data are shown as in Fig. 5. These simulations featured saltational transformation (Table 1).

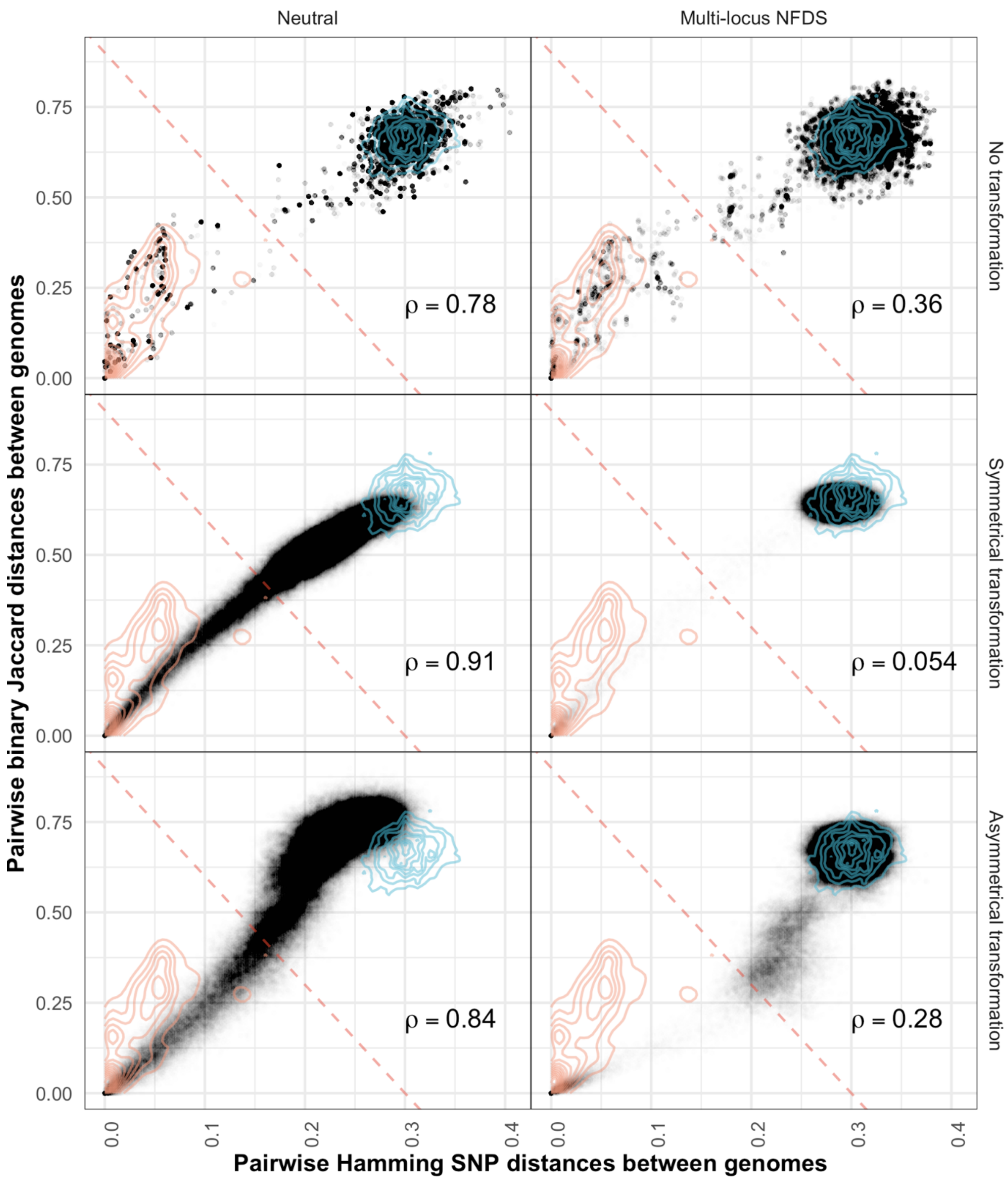


**Figure S14:** Scatterplots comparing the distributions of pairwise genetic distances between isolates at the final timepoint of simulations. Data are shown as in Fig. 4. These simulations featured inward migration (Table 1).

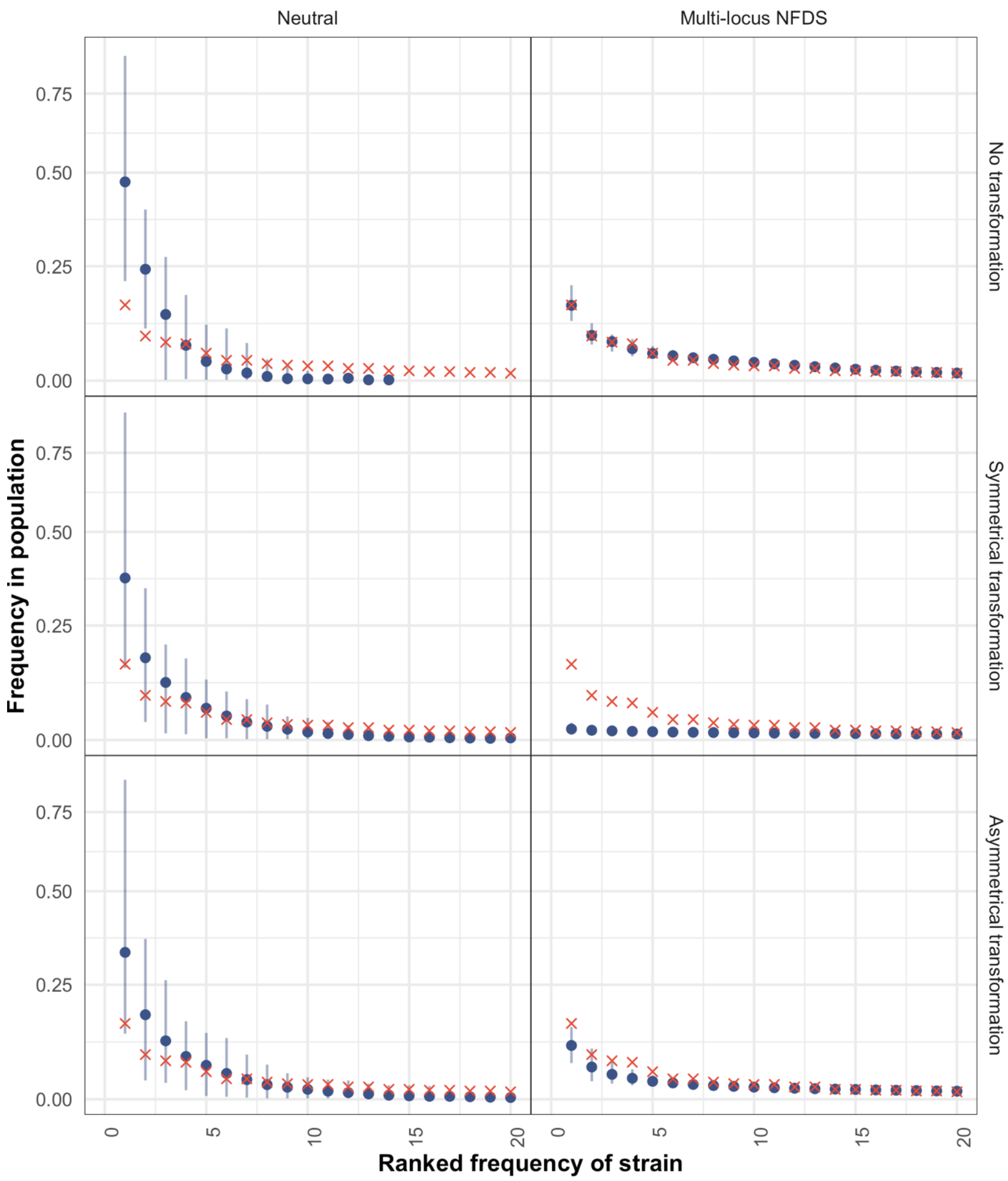


**Figure S15:** Scatterplots comparing the rank-frequency distributions of strains in the overall set of genomic data (red crosses), and those from samples of isolates from the final timepoint of 100 replicate simulations (blue points). Data are shown as in Fig. 5. These simulations featured inward migration (Table 1).

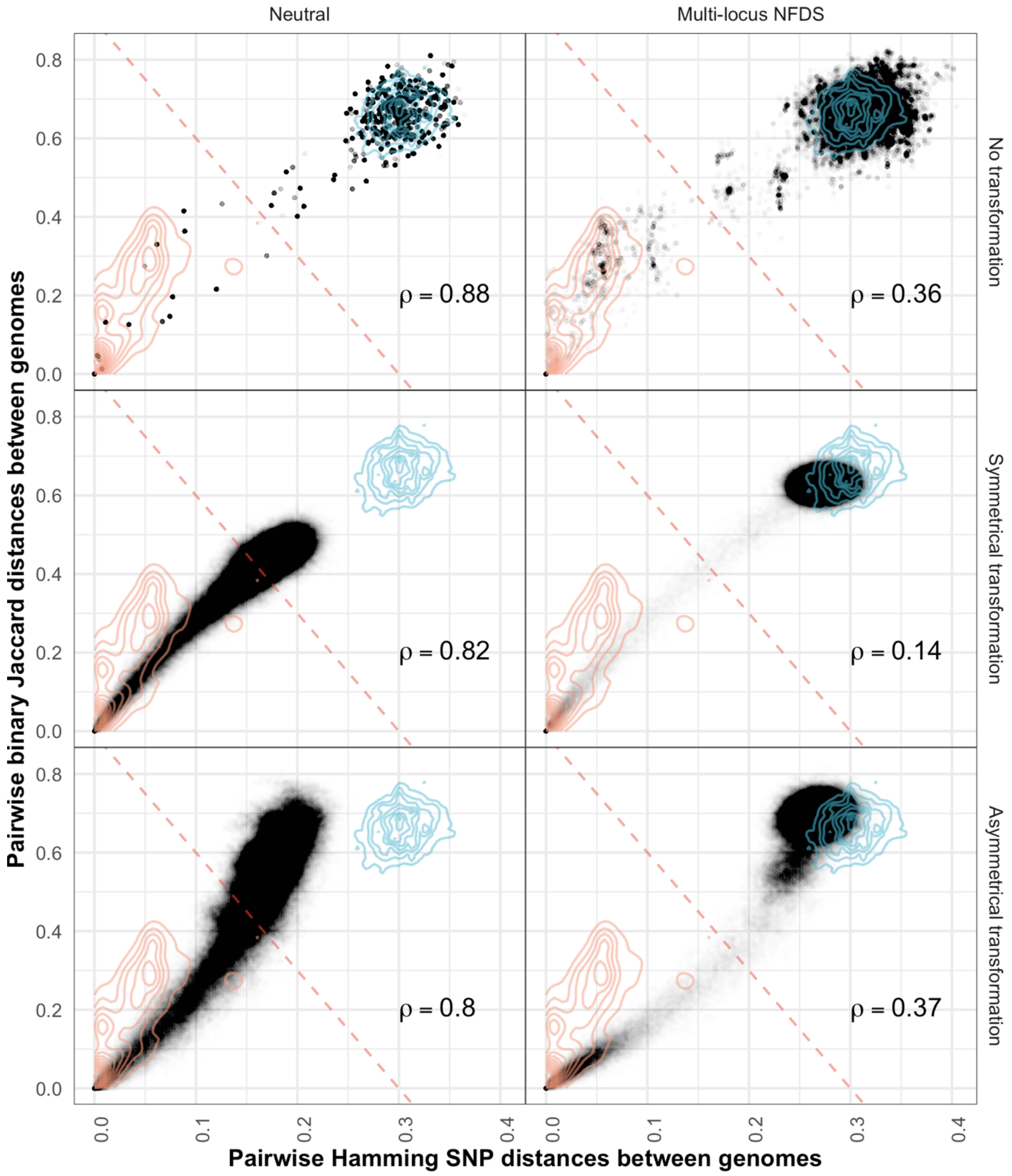


**Figure S16:** Scatterplots comparing the distributions of pairwise genetic distances between isolates at the final timepoint of simulations. Data are shown as in Fig. 4. These simulations featured weak multi-locus NFDS (Table 1).

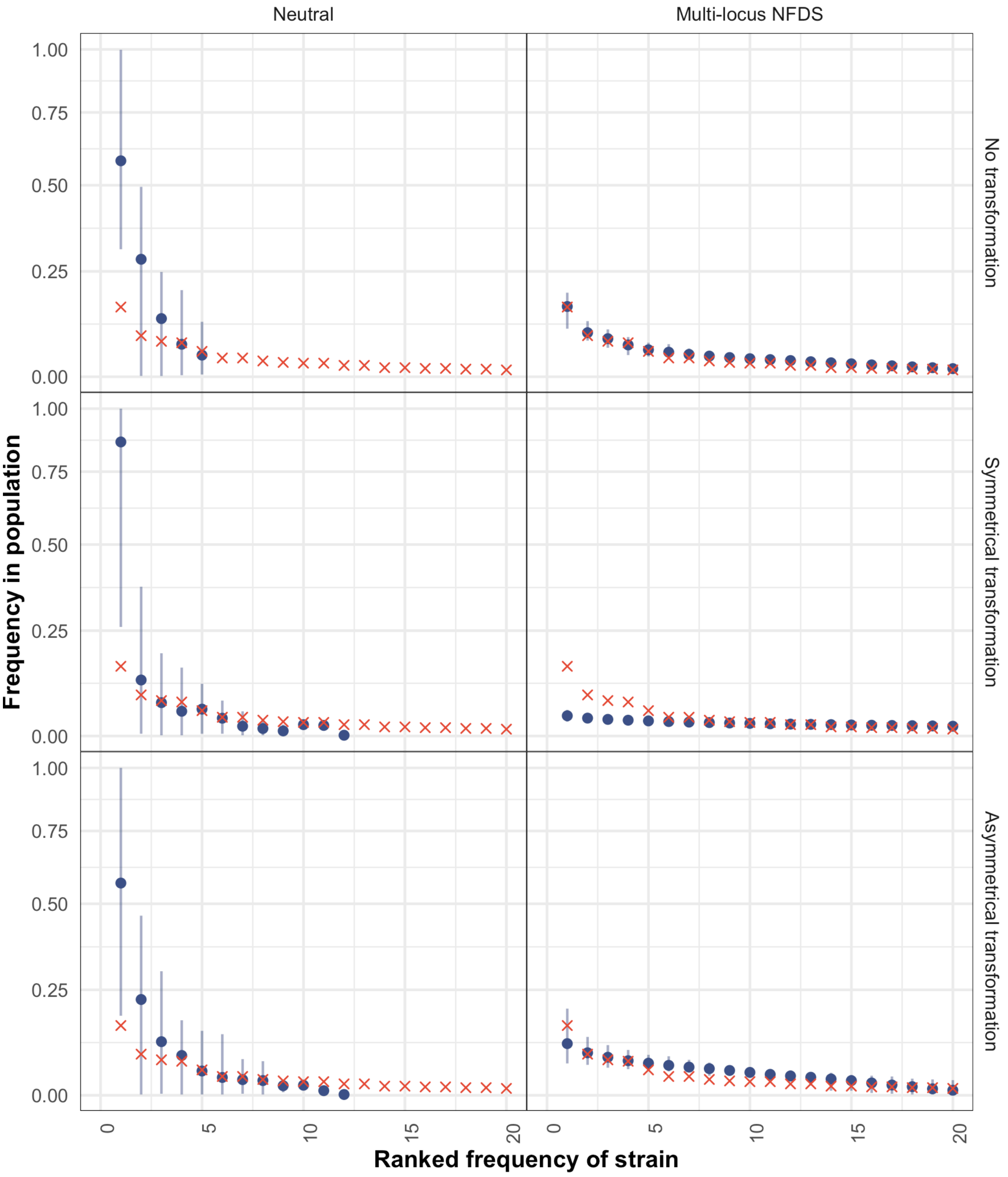


**Figure S17:** Scatterplots comparing the rank-frequency distributions of strains in the overall set of genomic data (red crosses) and those from samples of isolates from the final timepoint of 100 replicate simulations (blue points). Data are shown as in Fig. 5. These simulations featured weak multi-locus NFDS (Table 1).

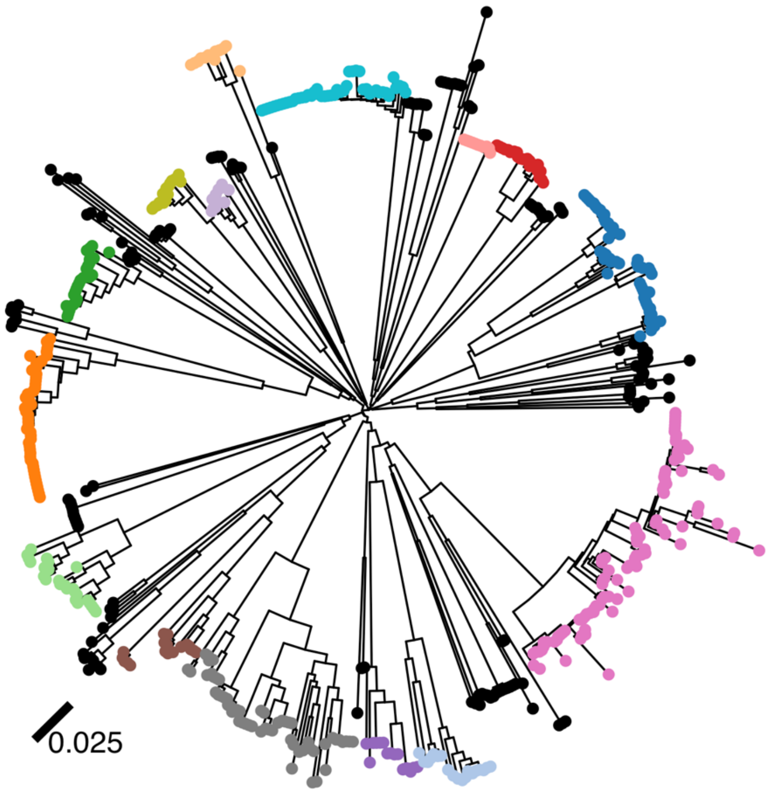


**Figure S18:** Neighbour-joining tree constructed from the intermediate-frequency core genome SNPs in the genomic data ($S$ = 1090). Tips corresponding to isolates belonging to common strains, with more than 10 representatives in the population, are coloured according to this categorisation; other tips, corresponding to isolates of rarer strains, are coloured black.

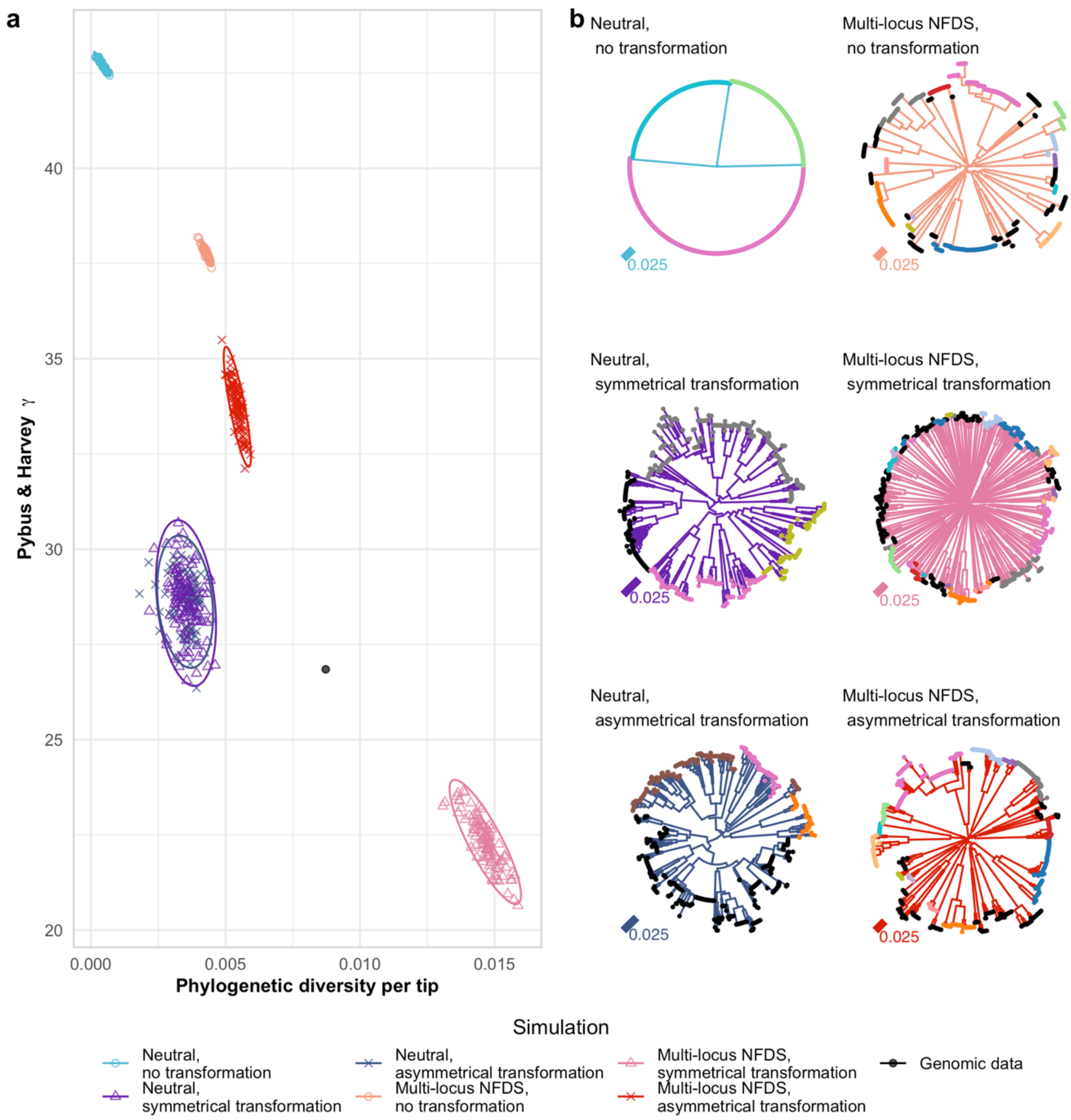


**Figure S19:** Comparison of trees between genomic data and simulation outputs. Data are shown as in Fig. 6. **a** Scatterplot comparing the characteristics of the neighbour-joining trees **b** Representative trees from individual simulations from each parameter set. These simulations featured saltational transformation (Table 1).

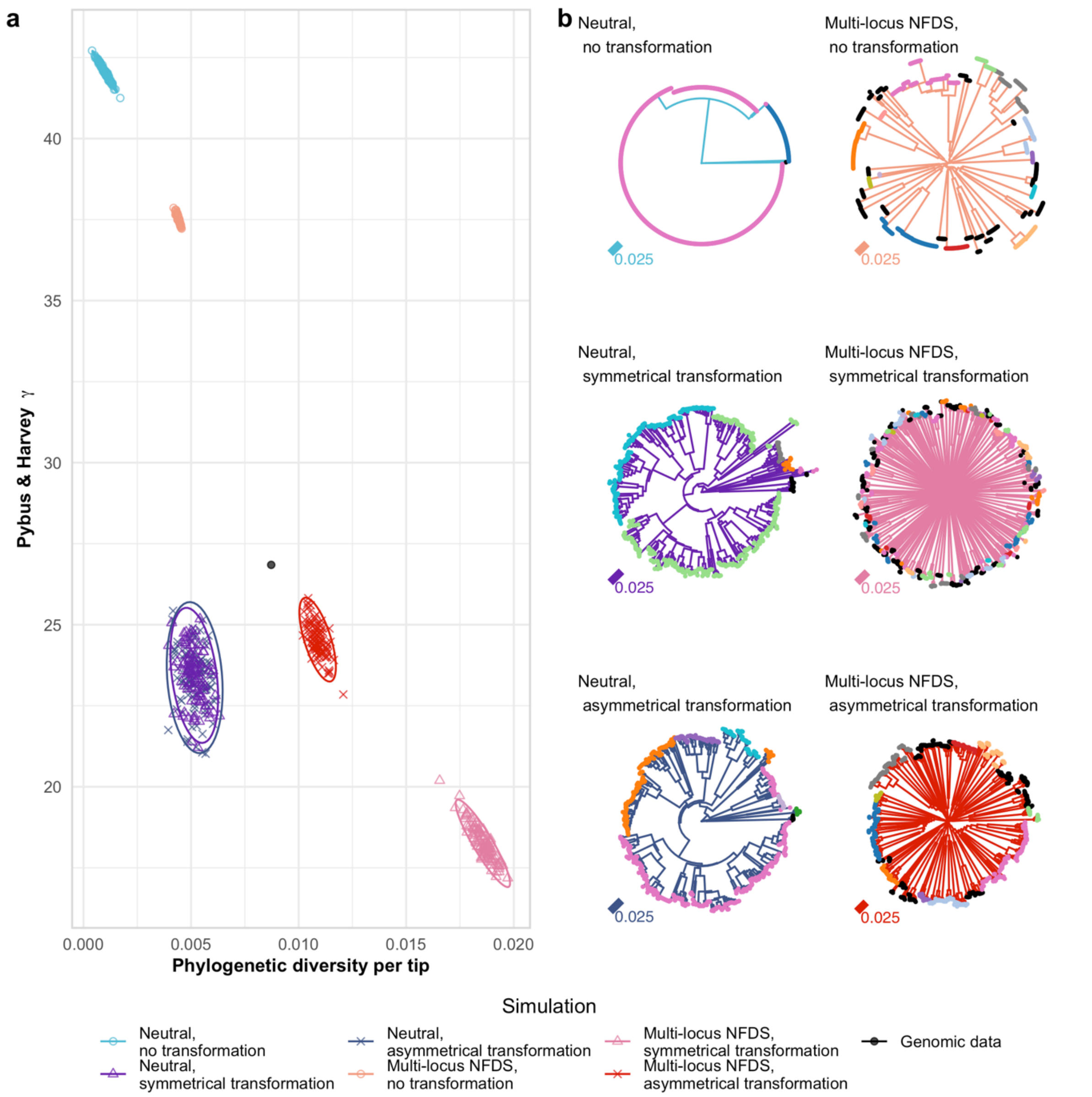


**Figure S20:** Comparison of trees between genomic data and simulation outputs. Data are shown as in Fig. 6. **a** Scatterplot comparing the characteristics of the neighbour-joining trees **b** Representative trees from individual simulations from each parameter set. These simulations featured inward migration (Table 1).

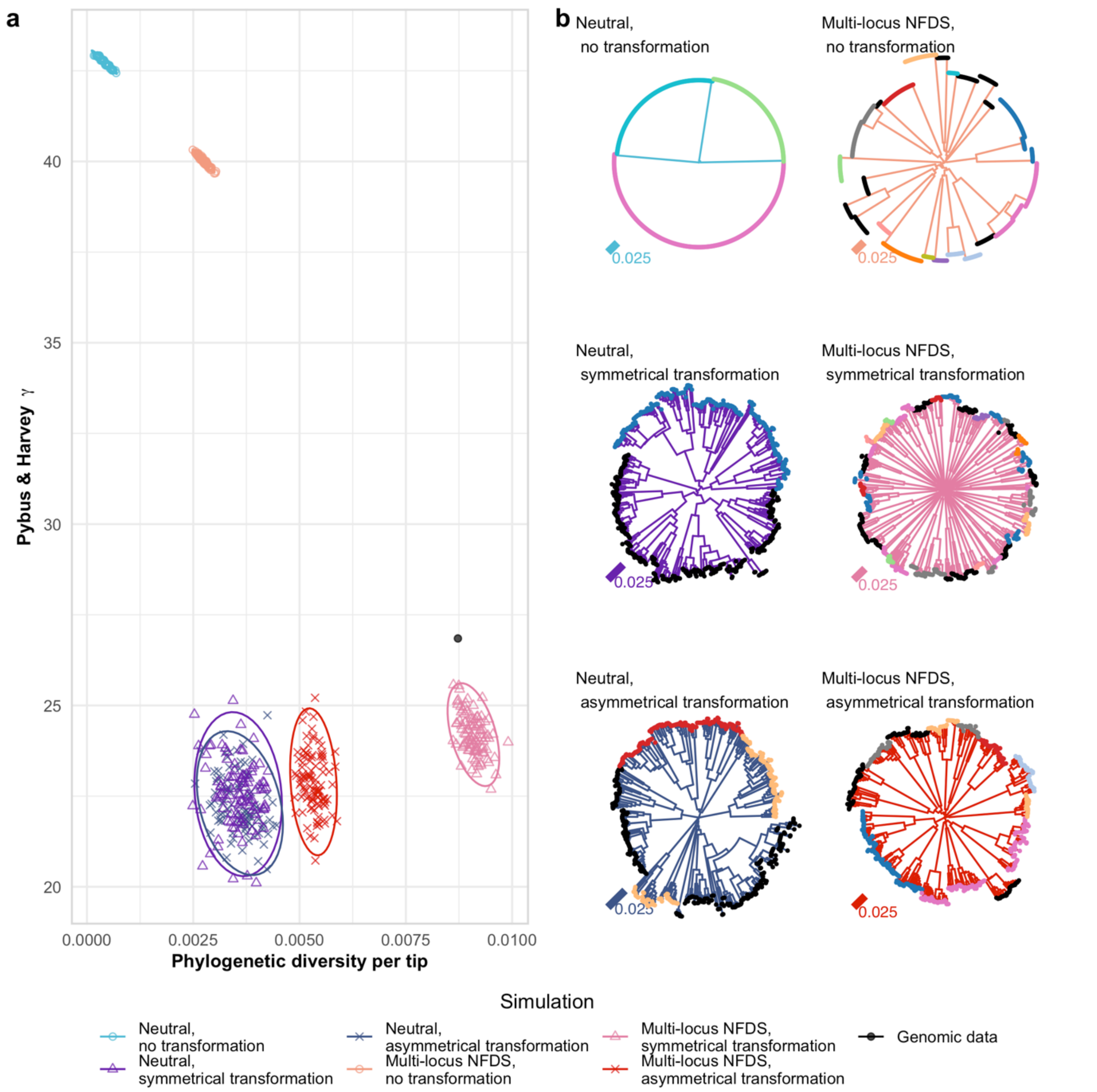


**Figure S21:** Comparison of trees between genomic data and simulation outputs. Data are shown as in Fig. 6. **a** Scatterplot comparing the characteristics of the neighbour-joining trees **b** Representative trees from individual simulations from each parameter set. These simulations featured weak multi-locus NFDS (Table 1).

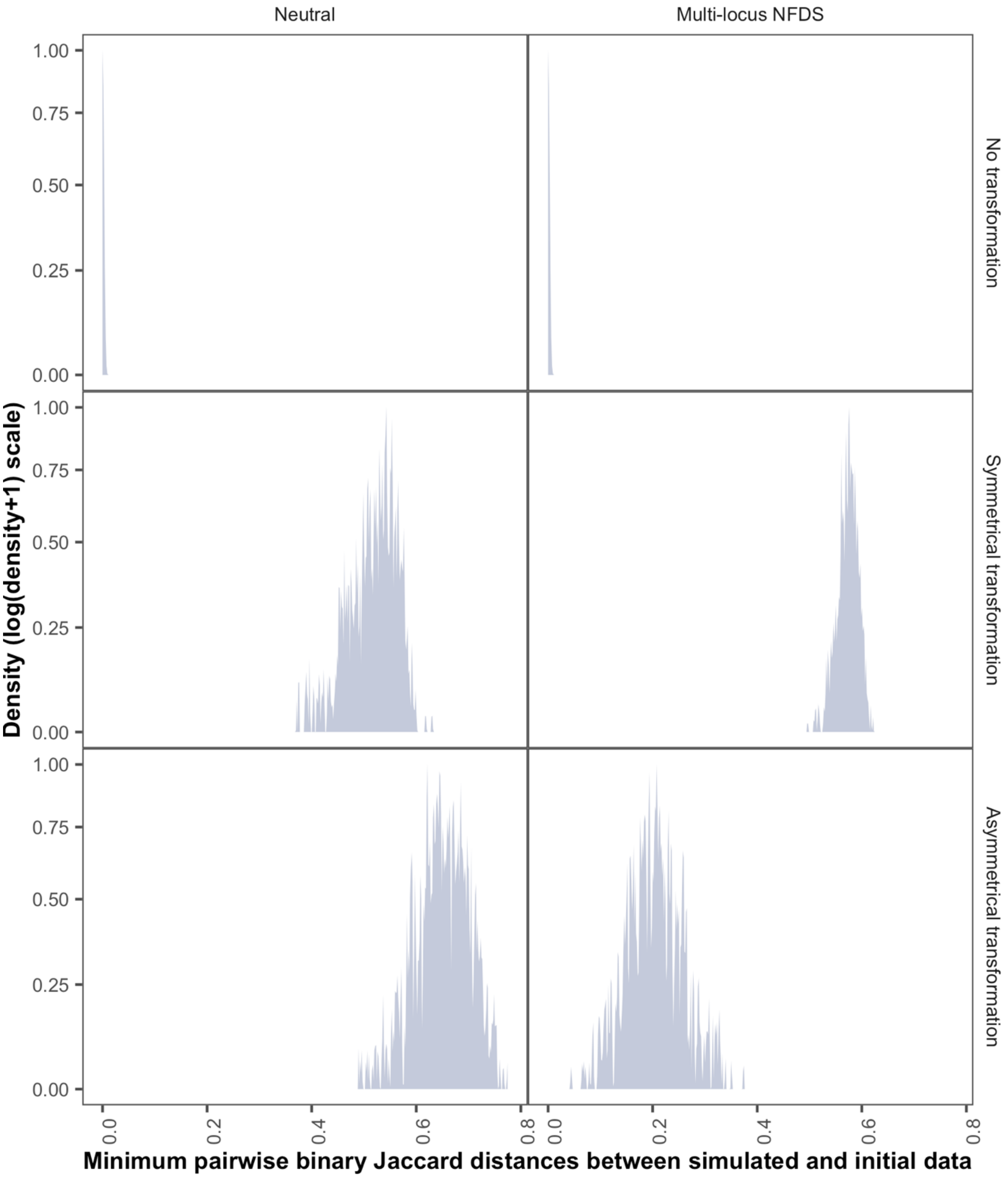


**Figure S22:** Density plot (using a bandwidth of 0.01) showing divergence of genotypes from the initial genomic data. The accessory locus content of each isolate sampled from the final timestep (*N* = 616) of each set of 100 replicate simulations was compared to the genomic data by calculating the pairwise binary Jaccard distances. These distributions show each isolate's minimum distance to a genotype in the genomic data (overall *N* = 61,600 per panel).

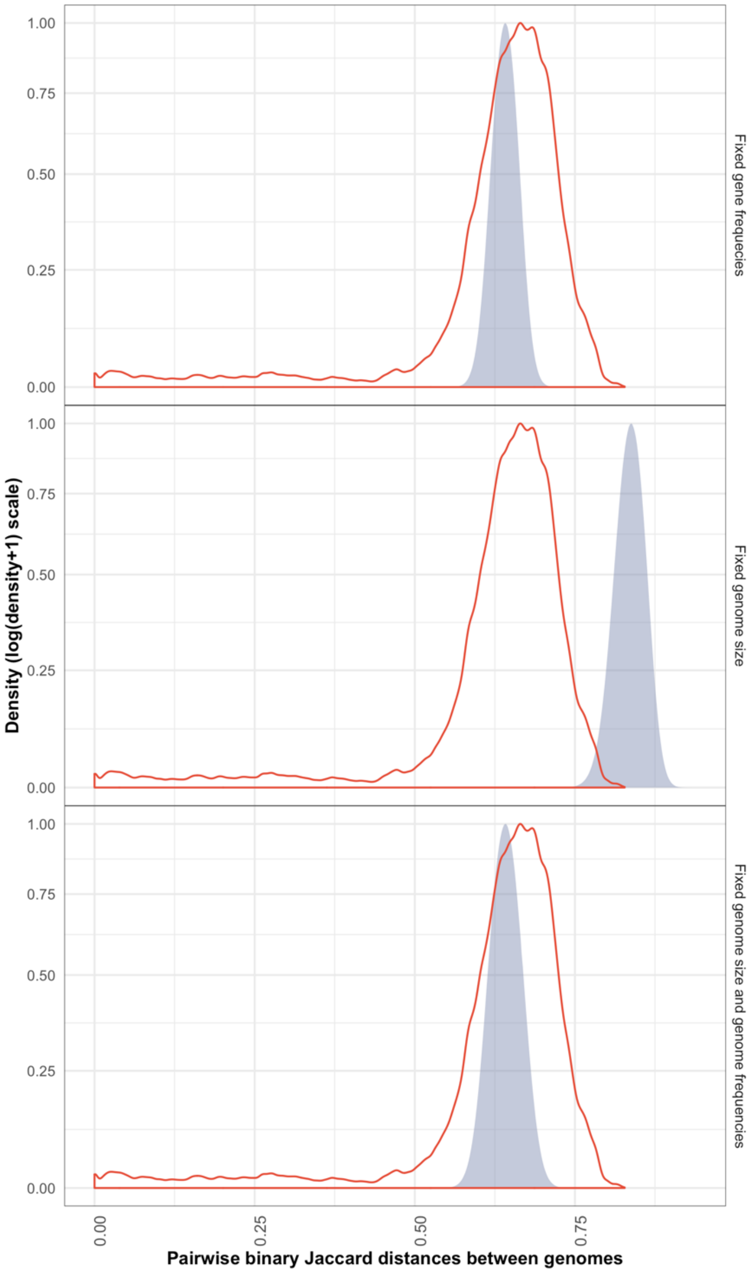


**Figure S23:** Density plot comparing the distributions of pairwise binary Jaccard distances, calculated from genotypes' accessory gene content, between isolates in the genome data (red line; $N$ = 379,456) and from genotypes generated by permuting the $g_{i,l}$ matrix 100 times (blue filled area; $N$ = 18,942,000 in each panel). The top plot shows the distances between genotypes generated by permuting the gene content while preserving genes at their equilibrium frequency. The middle plot shows the distances between genotypes generated by permuting the gene content while preserving the number of accessory genes in each genome. The bottom plot shows the distances between genotypes generated when permuting gene content under both constraints.

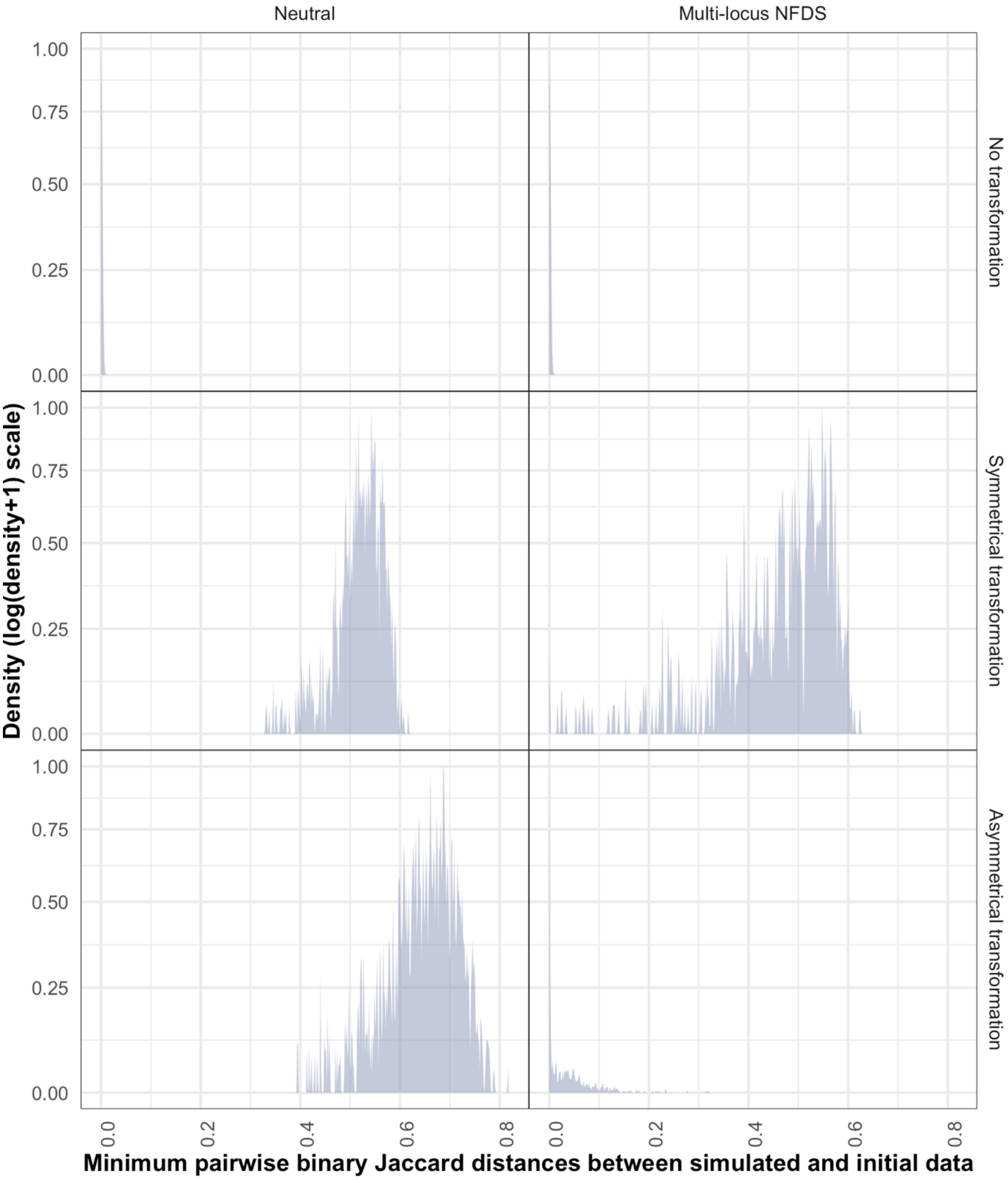


**Figure S24:** Density plot (using a bandwidth of 0.01) showing divergence of genotypes from the initial genomic data, as displayed in Fig. S22. These simulations featured saltational transformation (Table 1).
